# Distinct bile salt hydrolase substrate preferences dictate *C. difficile* pathogenesis

**DOI:** 10.1101/2022.03.24.485529

**Authors:** Matthew H. Foley, Morgan E. Walker, Allison K. Stewart, Sarah O’Flaherty, Emily C. Gentry, Garrison Allen, Shakshi Patel, Meichen Pan, Violet V. Beaty, Molly E. Vanhoy, Michael K. Dougherty, Sarah K. McGill, Ajay Gulati, Pieter C. Dorrestein, Erin S. Baker, Matthew R. Redinbo, Rodolphe Barrangou, Casey M. Theriot

**Affiliations:** Department of Pathobiology and Population Health, College of Veterinary Medicine, North Carolina State University, Raleigh, NC, USA; Department of Food, Bioprocessing and Nutrition Sciences, North Carolina State University, Raleigh, NC, USA; Department of Chemistry, University of North Carolina at Chapel Hill, Chapel Hill, NC, USA; Department of Chemistry, North Carolina State University, Raleigh, NC, USA; Skaggs School of Pharmacy and Pharmaceutical Sciences, University of California San Diego, La Jolla, San Diego, CA, USA; Collaborative Mass Spectrometry Innovation Center, Skaggs School of Pharmacy and Pharmaceutical Sciences, University of California San Diego, La Jolla, CA, USA; Department of Medicine, Division of Gastroenterology and Hepatology, University of North Carolina at Chapel Hill, Chapel Hill, NC, USA; Department of Pathology and Laboratory Medicine, University of North Carolina at Chapel Hill, Chapel Hill, NC, USA; Departments of Biochemistry & Biophysics, and Microbiology & Immunology, and the Integrated Program in Biological and Genome Sciences, University of North Carolina at Chapel Hill, Chapel Hill, NC, USA

**Keywords:** bile salt hydrolase, bile acid, live biotherapeutic, *Lactobacillus*, *Clostridioides difficile*

## Abstract

Bile acids (BAs) mediate the crosstalk between human and microbial cells and influence intestinal diseases including *Clostridioides difficile* infection (CDI). While bile salt hydrolases (BSHs) shape the BA pool by deconjugating conjugated BAs, the basis for their substrate preferences and impact on *C. difficile* remain elusive. Here, we survey the diversity of *Lactobacillus* BSHs and unravel the structural basis of their substrate preference. We show that leveraging BSH activity and specificity is an effective strategy to prevent *C. difficile* growth in clinically relevant CDI models. A range of non-canonical conjugated BAs is also identified, comprising unique BSH substrates that also inhibit *C. difficile* spore germination. These findings establish BSHs as intestinal enzymes essential to BA homeostasis and colonization resistance against *C. difficile*.

**One sentence summary:** Bile salt hydrolase activity inhibits *C. difficile* by shaping the conventional and non-canonical conjugated bile acid pools

## Introduction

Bile acids (BAs) are host-synthesized and microbial-derived metabolites that support intestinal health and homeostasis by providing a scaffold for host-microbiome crosstalk and adaptation (*1, 2*). Host-encoded BA receptors recognize BAs as signaling molecules to regulate key aspects of host immunity (*3*), metabolism (*4*), and circadian rhythms (*5, 6*). BAs also shape the microbiota, with specific taxa being inherently more resistant to bile stress by harboring mechanisms to survive BA exposure (*7, 8*). The influence of BAs on the microbiota composition is reciprocated by the microbial transformation of BAs to unlock the chemical complexity present in the intestinal BA pool (*9*). The resulting mosaicism of the BA pool is a hallmark of host health and is dependent on the collective activity of microbiome-encoded BA altering enzymes (*10*).

One such enzyme, bile salt hydrolase (BSH), comprises a family of microbial enzymes that catalyze a critical step in BA biosynthesis (*11*). Primary conjugated BAs synthesized in the liver, such as the tauro- and glyco-conjugates of cholic acid (TCA, GCA), are deconjugated by BSHs to generate cholic acid (CA) (**Fig. 1A**; BA names and abbreviations are summarized in **Fig. S1**). This vanguard catalytic step acts as a “gatekeeper” for subsequent BA transformations that turn primary BAs into secondary BAs (i.e. deoxycholic acid -DCA- formed from the 7α-dehydroxylation of CA) (*1, 11*).

**Figure 1.**
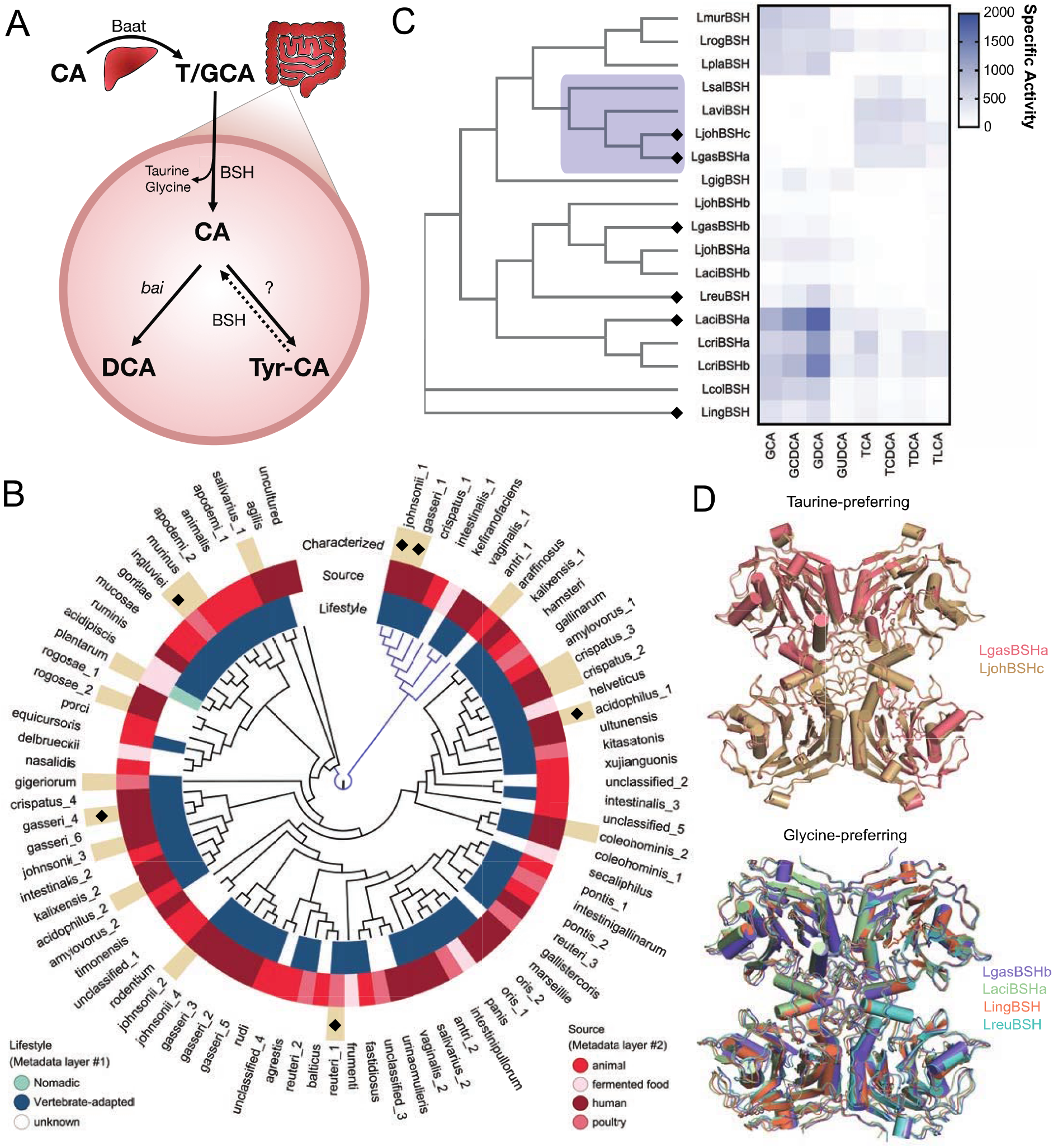
*Lactobacillus* BSHs have distinct substrate preferences. (A) Overview of BA metabolism using cholic acid (CA) as an example. BAs are synthesized and conjugated with a taurine or glycine (T/GCA) by the liver enzyme Baat. Conjugated BAs are deconjugated by BSHs, allowing transformations such as 7α-dehydroxylation encoded by the *bai* operon to generate secondary BAs like DCA. Deconjugated BAs can be re-conjugated to produce non-canonical conjugated BAs like Tyr-CA. (B) The *Lactobacillus* BSH phylogenetic tree was assembled from 84 BSH clusters. Major clades are distinguished by branch color. Each cluster name indicates the BSH-encoding species with the inner metadata layers indicating lifestyle and isolation source (*36*). The outer metadata layer indicates the BSHs cloned for characterization with diamonds indicating BSH crystal structures presented here. (C) Heatmap of BSH activity across conjugated BAs. The clade highlighted in blue displays preference for taurine-conjugated BAs. Values represent mean specific activity (n = 3-4). Phylogenetic relationships between BSHs are based on amino acid sequences. (D) Superposition of the taurine- and glycine-preferring BSH enzyme tetramer crystal structures presented here.

Dysregulation of BA biosynthesis and BSH activity has been directly implicated in obesity (*12*), cancer (*13*), inflammatory bowel disease (*3*), and colonization resistance against pathogens including *Clostridioides difficile* (*14, 15*). Strategically managing BSH activity to rationally toggle BA metabolism and judiciously alter intestinal pathobiology presents an emerging therapeutic approach. Additionally, the recent discovery of non-canonical conjugated BAs has considerably increased the complexity of the BA pool and altered the paradigm of BSH-BA biology. In particular, BSHs do not appear to process these non-canonical conjugated BAs *in vitro*, indicating that renewed and expanded investigations of BSH-substrate relationships are warranted (**Fig. 1A)** (*16–18*).

Here, we examine the phylogenetic, biochemical, structural diversity and specificity of BSHs in *Lactobacillus*, a prototypical genus for studying BA deconjugation. We define how the conjugated amino acid moiety drives substrate preferences in distinct BSHs and how those preferences can be leveraged to prevent *C. difficile* spore germination and colonization in pre-clinical models. We further reveal a new function for BSHs, the ability to process an expanded set of substrates including non-canonical conjugated BAs.

## Surveying *Lactobacillus* BSH Diversity

To map the range of BSHs harbored by *Lactobacillus*, 3,712 genomes from 274 lactobacilli were assembled to comprehensively define the diversity of the enzyme within this key intestinal genus (**Table S1)** (*19*). Highly similar BSH sequence (>95% amino acid identity) were grouped into 84 clusters and a representative sequence from each cluster was used to construct a phylogenetic tree. Nearly all BSH clusters are identified in vertebrate-adapted species and ~40% were human-associated, underscoring their role in host-microbiota symbiosis (**Fig. 1B, Table S1**).

To move toward a functional understanding of BSH activity *in vivo*, 17 diverse BSHs were selected for heterologous expression and purification **(Table S2, Fig. S2)**. BSH activity was screened on a panel of conjugated primary and secondary BAs to comprehensively determine BSH substrate preferences **(Fig. 1C)**. BSHs display a well-defined bias for either glycine or taurine-conjugated BAs. Glycine-preferences dominate the BSHs tested, potentially reflecting the increased abundance of glycine-conjugated BAs among vertebrates (*20*), whereas taurine-specificity is restricted to only a few related BSHs. The glycine-preferring clade containing LaciBSHa, LcriBSHa, and LcriBSHb exhibits the highest activity. Furthermore, the taurine-preferring LaviBSH, LjohBSHc, and LgasBSHa (corresponding to the clusters arrafinosis, johnsonii_1, and gasseri_1, respectively) enzymes are all present within a single major clade while the LsalBSH (cluster salivarius_2) stands alone in a distantly related branch. This suggests that taurine-preference may have evolved twice within *Lactobacillus*. Overall, BSH specialization for glycine or taurine illustrates how lactobacilli may have tailored their metabolism to manage BA exposure.

## Structural Basis for BSH Substrate Preference

To dissect the molecular basis of BSH substrate specificity, we determined the crystal structures of six of the *Lactobacillus* BSHs examined above, four glycine-preferring (LgasBSHb, LaciBSHa, LingBSH, LreuBSH) and two taurine-preferring BSHs (LgasBSHa, LjohBSHc; **Fig. 1D, S3A, Table S3**). While most structures were unliganded, LgasBSHa was resolved in complexes with taurine or taurine plus CDCA. All BSHs form tetramers and each monomer exhibits the core four-layered αββα fold typical of the Ntn-hydrolase family (*21*), with RMSD values ranging from 0.5 to 2.0 Å over equivalent Cα positions (**Fig. 2A, S3B**). Within this standard core structure, we noted that each monomer contains a loop donated from a neighboring monomer within the tetramer (**Fig. 2A**). Importantly, sequence analysis revealed that taurine-preferring enzymes contained an S-R-G/S motif in this loop region, while the glycine-preferring enzymes encoded a G-V/T-G motif (**Fig. 2B, Fig. S4**) (*22*). These particular amino acids were also placed directly in each enzyme’s active site within the functional BSH tetramers (**Fig. 2C, 2D**). Thus, we hypothesized that these 3-residue motifs play important roles in BSH substrate preferences.

**Figure 2.**
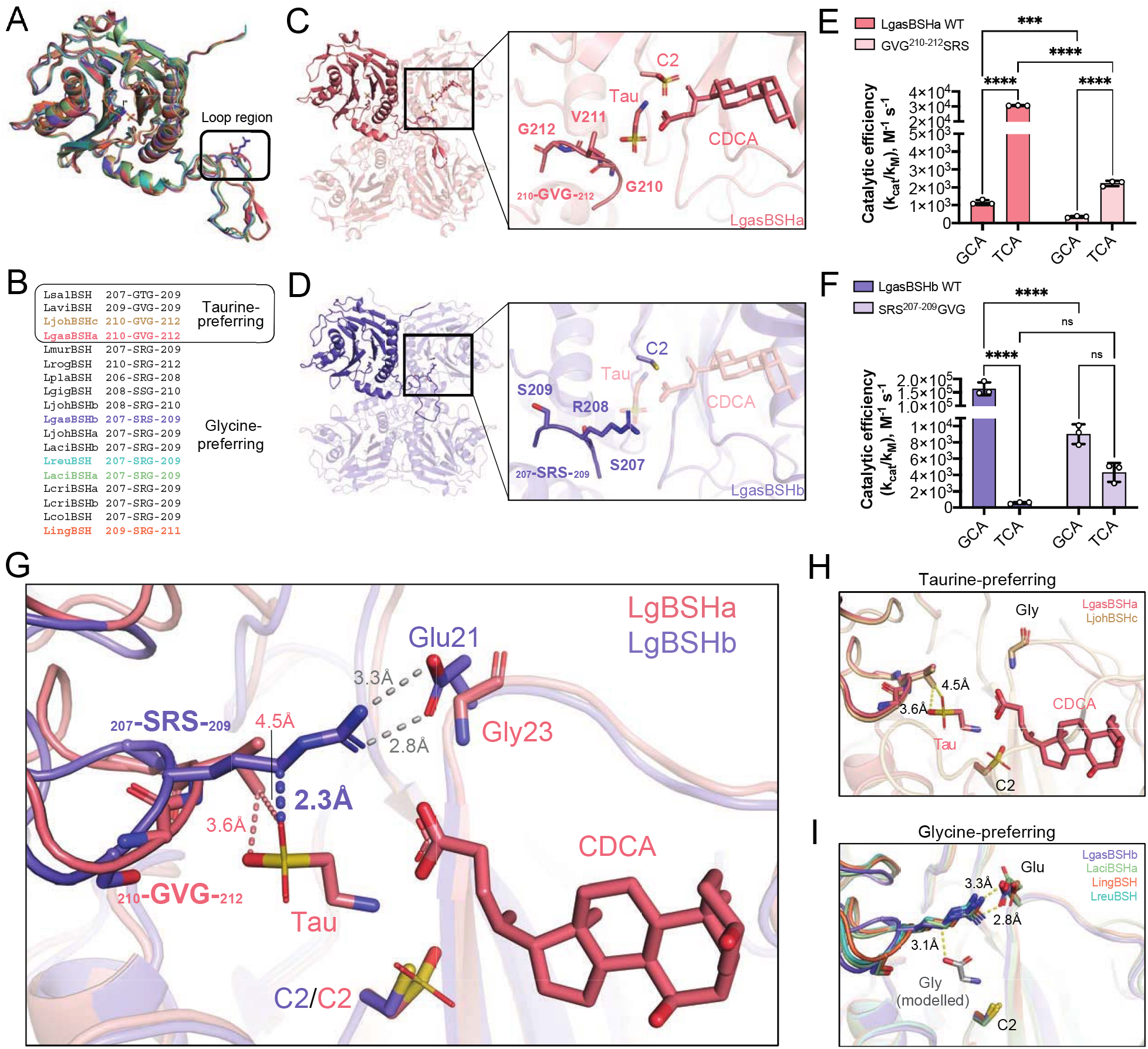
BSH loop region dictates enzyme substrate preference. (A) Superimposed monomers of the six BSH monomer structures presented here highlighting the substrate preference loop and colored as indicated in panel B. (B) Sequence alignment of the loop region of *Lactobacillus* BSHs. Resolved structures are colored as indicated, with a constant coloring scheme throughout the figure. Taurine-preferring enzymes exhibit a G-V/T-G motif, while glycine-preferring enzymes maintain a S-R-G/S motif. (C) The LgasBSHa tetramer with the active site shown in the inset, with the catalytic cysteine at position 2, the bound taurine and CDCA compounds, and the GVG swapped into the active site from the neighboring monomer highlighted. (D) The LgasBSHb tetramer with the active site shown in the inset, with the catalytic cysteine at position 2 and the SRS swapped into the active site from the neighboring monomer highlighted. The taurine and CDCA ligands from the LgasBSHa structure are also rendered transparently. (E, F) Catalytic efficiency for wild-type and mutant forms of LgasBSHa and LgasBSHb with GCA and TCA. Bars are the average of n=3 ± standard deviation. Significant differences were tested by two-way ANOVA with Tukey’s multiple comparisons test (ns not significant, \*\*\**p* < 0.001, \*\*\*\**p* < 0.0001). (G) Superposition of the active sites of LgasBSHa and LgasBSHb showing that taurine is accommodated by Val211 of LgasBSHa but clashes at 2.3 Å with the side chain of Arg208 in LgasBSHb, which is held in place via a salt bridge with Glu21. (H, I) Superpositions of the active sites of the two taurine-preferring or the four glycine-preferring BSH structures presented here reveals consistent active site architectures that accommodate taurine or glycine, respectively.

To test this, the 210-GVG-212 region of taurine-preferring LgasBSHa was replaced with SRS, while the 207-SRS-209 region of glycine-preferring LgasBSHb was replaced with GVG, and catalytic efficiencies with conjugates of CA were examined. The LgasBSHa-SRS variant enzyme exhibited significantly reduced catalytic efficiency with TCA, the wild-type enzyme’s preferred substrate, while also reducing its processing of GCA (**Fig. 2E**). Similarly, the LgasBSHb-GVG variant showed significantly lower catalytic efficiency with its preferred GCA substrate, and a trend toward increased activity with the non-standard substrate TCA (**Fig. 2F**). Taken together, these functional data support the structure-guided hypothesis that short sequence motifs play important roles in defining the marked glycine *vs*. taurine substrate preferences exhibited by *Lactobacillus* BSHs, reflecting specialization.

Structurally, the presence of R208 in LgasBSHb appears to block the processing of taurine-conjugated BAs. When the LgasBSHa and LgasBSHb active sites are overlaid, the distance between LgasBSHb R208 and taurine from the LgasBSHa structure is 2.3 Å, which would generate a steric clash (**Fig. 2G**). In contrast, V211 in LgasBSHa is 3.6 Å away and would not occlude the binding of taurine-conjugated BAs (**Fig. 2G**). Furthermore, R208 in LgasBSHb forms a hydrogen bond with E21 and that interaction creates a more enclosed active site than that seen in taurine preferring BSHs (Fig. 2G). In support of this conclusion, the active sites of the four additional BSH structures resolved here (LjohBSHc, LaciBSHa, LingBSH, LreuBSH) reveal analogous architectures (**Fig. 1D, 2H, I**). Importantly, when a glycine conjugate was modelled into the active site of the glycine-preferring BSHs, the distance between R208 and glycine increased to 3.1 Å, indicating that this smaller substrate can be accommodated with this clade of enzymes (**Fig. 2I**). Together, these data define three-residue motifs as crucial for the glycine vs. taurine conjugate preferences demonstrated by *Lactobacillus* BSHs.

## Bile Acids Inhibit *C. difficile* in a Conjugation-Dependent Manner

*C. difficile* infection (CDI) is a significant public health problem that can be difficult to treat with conventional antibiotics (*23*). Even after successful treatment with vancomycin, there remains a high recurrence rate (~30%) (*24*), making a fecal microbiota transplant (FMT) the last line of treatment (*25*). Although FMTs are a lifeline for recurrent CDI (rCDI) patients, the long-term consequences are unknown at this time, making a targeted therapeutic approach urgently needed. Susceptibility to CDI is predominantly driven by antibiotic-driven perturbations to the intestinal microbiota, which results in a loss of colonization resistance (*26, 27*). Strong evidence suggests that microbial BA metabolism is an important mechanism of colonization resistance against *C. difficile* (*28*), as antibiotic usage causes depleted BSHs and increases conjugated BAs (*14*).

BAs govern fundamental aspects of *C. difficile*’s pathogenesis: TCA acts as a germinant for *C. difficile* spores (*29*), whereas CDCA can inhibit germination and growth (*30*). Most studies have focused on the production of deconjugated secondary BAs as inhibitors of *C. difficile* pathogenesis while few have examined the contributions of BSH activity (*31*). The therapeutic potential of BSHs has been proposed, but gaps in our knowledge of their pleiotropic effects on intestinal biology and their exact mechanism of action have hampered their development (*32, 33*).

To determine how BA deconjugation impacts *C. difficile*, we measured the ability of common BAs to inhibit germination in the presence of TCA (**Fig. 3A-C**). Though all BAs tested inhibit germination compared to TCA alone, deconjugated BAs are significantly more inhibitory compared to their taurine-conjugated versions, whereas they are equally or only mildly more inhibitory relative to their glycine-conjugated variants (**Fig 3B**). We also assayed the ability of BAs to inhibit *C. difficile* growth, as we hypothesized that the conjugation would determine BA toxicity (**Fig. 3C**). Taurine-conjugates inhibit growth less while glycine-conjugates and deconjugated BAs are much more toxic. Moreover, taurine-conjugated BAs are significantly less disruptive to *C. difficile’* membrane integrity compared to glycine-conjugated and deconjugated BAs (**Fig. S5**). Overall, using BSHs to deconjugate taurine-conjugated BAs may be a potent strategy to restrict *C. difficile* in the intestinal tract.

**Figure 3.**
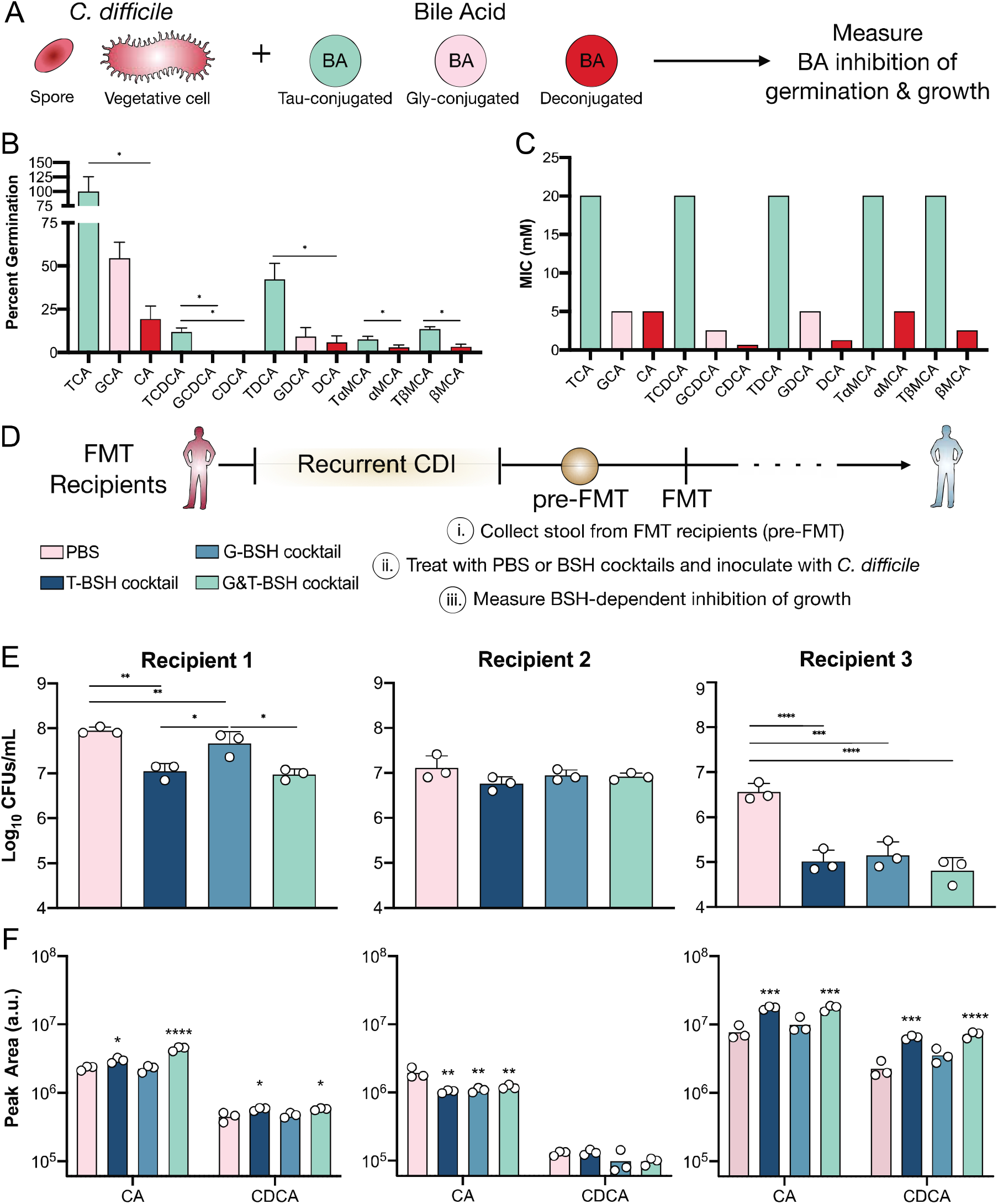
Deconjugated BAs and BSH activity can inhibit *C. difficile*. (A) Schematic of *in vitro* BA-dependent inhibition of *C. difficile* germination and growth. (B) Inhibition of spore germination using various BAs (0.75 mM) with the germinant TCA (2 mM). (C) Inhibition of *C. difficile* growth as determined by minimum inhibitory concentration (MIC) testing. Statistical differences in B and C were assessed by a one-tailed Welch’s *t* test. (D) Schematic of pre-FMT collection and *ex vivo* growth. Pre-FMT stool collected from rCDI patient was supplemented with PBS or a BSH cocktail and subsequently inoculated with *C. difficile*. (E) *C. difficile* growth was measured at 8h. (F) Targeted metabolomics showing the deconjugated BAs CA and CDCA from the samples from E. Statistical differences tested by one-way repeated measures ANOVA with Sidak’s correction for multiple comparisons. All bars represent the average of n=3 replicates ± sd. \**p* < 0.05, \*\**p* < 0.01, \*\*\**p* < 0.001, \*\*\*\**p* < 0.0001.

To determine whether BSHs are sufficient to inhibit *C. difficile* and provide an alternative approach to using antibiotics or FMT, stool from three rCDI patients was collected prior to them receiving an FMT (**Fig 3D**). Unlike mice, FMT recipients have a complex mixture of taurine and glycine-conjugated primary BAs (**Fig S6**). To test whether BSH substrate preference could impact *C. difficile* growth, pre-FMT stool was used to culture *C. difficile* and samples were treated with PBS, a cocktail of taurine-preferring BSHs (T-BSH), glycine-preferring BSHs (G-BSH), or a broadly-acting combination (G&T-BSH) **(Fig. 3E, S7)**. Alone, the BSH cocktails, glycine, or taurine have no impact on *C. difficile* growth (**Fig. S8**). When Recipient 1’s stool is treated with the T-BSH and G&T-BSH cocktails, *C. difficile* growth is significantly inhibited at 8 h. Recipient 2’s stool doesn’t alter *C. difficile* growth when treated with any BSH cocktail, though the T-BSH and G&T-BSH cocktails slightly suppress growth at 24 h. Recipient 3’s stool robustly suppresses *C. difficile* growth at 8 h using all BSH cocktails, but only the T-BSH and G&T-BSH cocktails maintain that suppression at 24 h.

Targeted BA metabolomics was used to assess how the BSH cocktails alter the pre-FMT BA pool and explain the individual variability between recipients (**Fig 3F, S9**). Recipients 1 and 3 display an increase in CA and CDCA primarily in response to the T-BSH and G&T-BSH cocktails. Recipient 3’s high levels of conjugated BAs prior to BSH treatment explains the BSH cocktail’s robust inhibition (**Fig. S6, 9**), Furthermore, Recipient 2 does not display an increase in deconjugated BAs from any BSH treatment, likely due to the overall low concentration of conjugated BAs prior to BSH treatment (**Fig S6**), thereby explaining why no *C. difficile* inhibition was observed in this sample.

## BSH Activity Inhibits *C. difficile* Spore Germination and Growth

To interrogate the role of BSHs in colonization resistance against *C. difficile*, we leveraged an antibiotic treated mouse model of CDI, since mice typically have a relatively rich amount of taurine conjugated bile acids. *C. difficile* initiates its infectious cycle when spores arrive and germinate in the small intestine, and then begin to grow and progress to the large intestine where the majority of toxin production and disease occurs. To examine the contribution of BSH activity to colonization resistance along the intestinal tract, we collected murine small intestinal and cecal contents from cefoperazone-treated mice at several times post-antibiotic treatment and supplemented them with PBS or the G&T-BSH cocktail (**Fig. 4A)**. The treated contents were then inoculated with 10^5^ spores or CFUs/mL to measure *ex vivo* spore germination and growth, respectively. Remarkably, the addition of the BSH cocktail significantly inhibits *C. difficile* spore germination in the small intestinal contents of all mice and suppresses growth at 8 and 24 h (**Fig. 4B, C, S10)**. Inhibition of growth is more subtle in the cecum, likely due to the lower concentration of conjugated BAs in this region of the intestinal tract (**Fig. 4D, S11**). Targeted metabolomics was used to corroborate the BSH cocktail’s effect on the murine BA pool. Small intestinal samples from day 0 treated with the BSH cocktail display a significant increase in deconjugated BAs (CA, CDCA, βMCA, and UDCA) and a decrease in the taurine-conjugated forms of the same BAs. In the cecum, only CA was significantly deconjugated (**Fig. 4D, S11**). By day 7 post antibiotic, some mice displayed higher amounts of deconjugated BAs, independent of BSH treatment, coinciding with the return of colonization resistance after antibiotic treatment.

**Figure 4.**
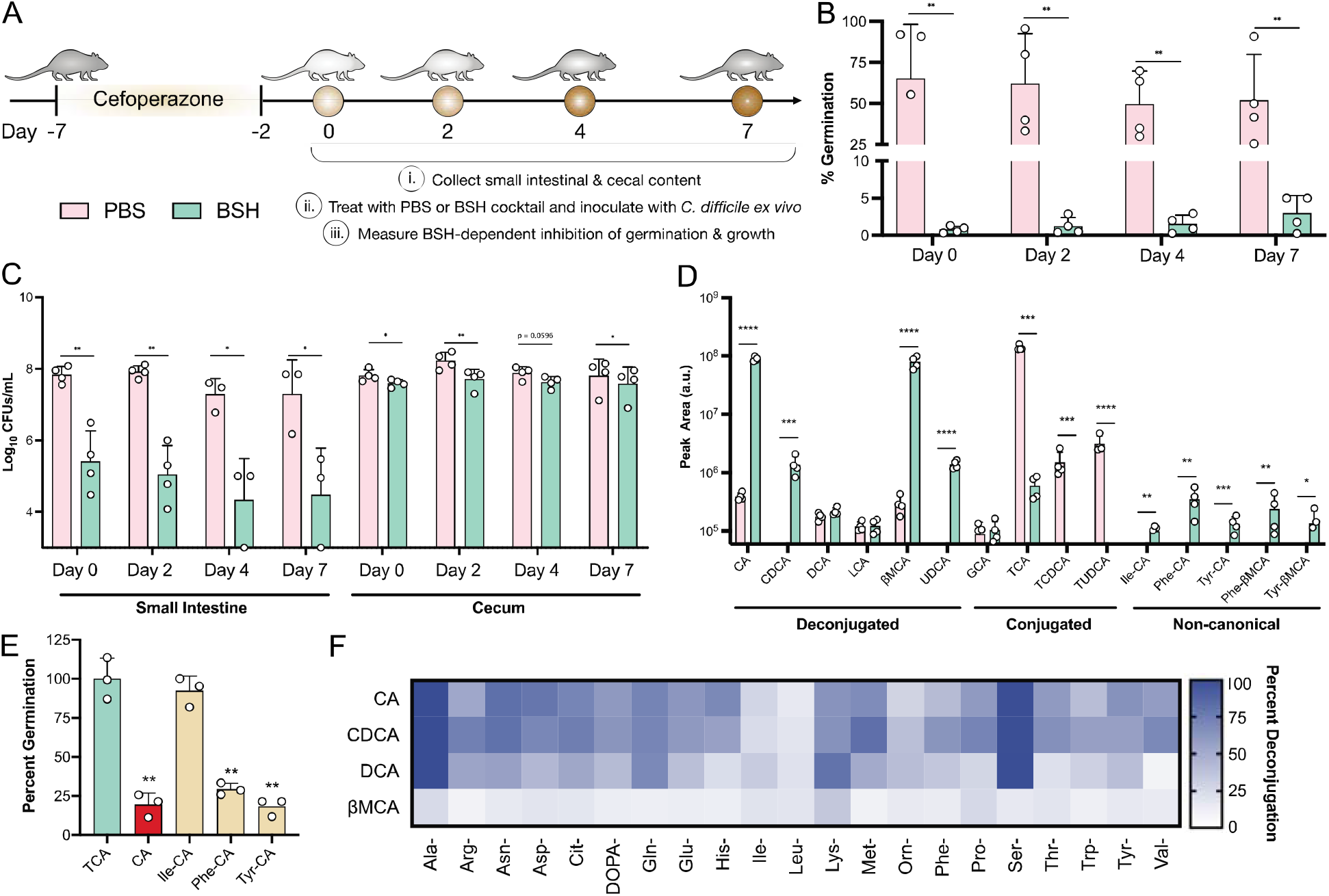
BSHs shape the non-canonical conjugated BA pool that governs *C. difficile* spore germination. (A) Schematic of BSH-dependent inhibition of *ex vivo C. difficile* germination and growth. Intestinal contents harvested from cefoperazone-treated mice (n = 3-4) were supplemented with a cocktail of BSHs and then inoculated with *C. difficile* spores or cells to measure germination or growth, respectively. (B) Germination of *C. difficile* spores in murine small intestinal contents. Contents were treated for 8h with PBS or a BSH cocktail before inoculation. (C) Growth at 8h of *C. difficile* in murine intestinal contents. Contents were treated with PBS or a BSH cocktail and inoculated simultaneously. Statistical differences were tested by ratio paired *t* test. (D) Targeted metabolomics showing the BAs identified in the Day 0 small intestinal samples from C. (E) Inhibition of *C. difficile* germination using various BAs (0.75 mM) with the germinant TCA (2 mM). Statistical differences were assessed by one-tailed *t* test. Bars represent mean ± standard deviation. (F) Heatmap of LingBSH across all non-canonical BA pools. Values represent the mean from n = 3 replicates of % deconjugation after 5 minutes. Bars represent mean of n=3-4 ± standard deviation. \**p* < 0.05, \*\**p* < 0.01, \*\*\**p* < 0.001, \*\*\*\**p* < 0.0001.

Aside from the increase in deconjugated BAs, the most striking difference in the BA pool is the unexpected increase in non-canonical conjugated BAs in the BSH-treated small intestinal samples (**Fig 4D, S11**). An array of intestinal bacteria can catalyze the conjugation of BAs to a variety of amino acids through an unknown mechanism (**Fig. 1A**) (*16, 17*). Despite their widespread synthesis, there is no evidence at this time that shows that the non-canonical conjugated BAs Ile/Leu-CA, Phe-CA, and Tyr-CA can be deconjugated (*16*). Ile-CA, Phe-CA, and Tyr-CA were among the BAs detected in the small intestinal samples. Given the increased CA and βMCA in the BSH-treated samples, these BAs are likely re-conjugated by the small intestinal microbiota to produce the non-canonical BAs observed. While these BAs did not affect *C. difficile* growth or induce spore germination (**Fig. S12A, B**), Tyr-CA and Phe-CA are able to inhibit spore germination, suggesting for the first time a role for the non-canonical conjugated BAs in colonization resistance against *C. difficile* (**Fig. 4E, S12C**).

## BSHs Process Non-Canonical Bile Acids Conjugates

To determine whether BSHs are able to process non-canonical conjugated BAs, we screened a larger range of BSHs and BAs (**Fig. S13**). While activity with these non-canonical conjugates was low compared to the canonical conjugations, some BSHs, like LingBSH and LreuBSH, exhibit appreciable activity on non-canonical BAs. To understand the recalcitrance of non-canonical conjugated BAs, we broadly monitored the deconjugation of 5 BSHs (LingBSH, LgasBSHa, LgasBSHb, LjohBSHc, and LreuBSH) with assorted preferences (**Fig. S14A**). BSHs were incubated with non-canonical conjugated BAs pools (AA-CA, AA-CDCA, AA-DCA, and AA-βMCA) that consist of one BA (CA, CDCA, DCA, or βMCA) conjugated to 22 different amino acids, excluding taurine and glycine (*18*), and initial BA deconjugation was quantified using LC-IMS-MS (**Fig. S14B**). LingBSH displays exceptional activity, suggesting that some BSHs may be uniquely adapted to process this expanded repertoire of BAs (**Fig. 4F, 14B, S15**). Ala and Ser-conjugated BAs are efficiently processed, and these observations were validated using purified Ala-CA and Ser-CA demonstrating that their susceptibility to deconjugation is similar to GCA and TCA (**Fig. S13**). Ala and Ser are the smallest amino acids next to Gly, suggesting that the steric constraints of the active site could dictate non-canonical conjugate deconjugation (**Fig. S15, S16)**. Notably, amine-containing amino acids such as Lys, Gln, and Asn are enriched and among the most efficiently processed conjugates (**Fig. S16**).

We also examine whether non-canonical conjugated BA processing would occur in BSH-expressing *L. gasseri* cells. Wild-type, Δ*bshA*, Δ*bshB*, and Δ*bshAB L. gasseri* was grown and reacted with the AA-CDCA pool and supernatants were used to detect deconjugation. Δ*bshAB* displayed some deconjugation, possibly due to cells sequestering BAs (**Fig. S14C)**. Individual BSH activities and substrate preferences were comparably maintained *in vivo*, demonstrating that BSH-expressing bacteria are capable of non-canonical BA deconjugation, and that they do so based on the enzymatic preferences of the BSHs they encode.

## Discussion

There is a growing appreciation for the roles that BSH substrate preferences play in the management of disease (*6, 8*). This work stresses the distinction between the deconjugation of glycine and taurine-conjugated BAs, the latter of which has been shown to play a role in the establishment of colonization resistance against *Klebsiella pneumoniae* and the development of colorectal cancer (*34, 35*). Our analysis of *C. difficile-BA* sensitivities motivated the use of glycine and taurine-preferring *Lactobacillus* BSHs to optimally inhibit *C. difficile ex vivo* (**Fig. 3D**). Thus, leveraging BA metabolism to restore colonization resistance may be an alternative therapy that avoids antibiotic damage, the unspecified risks of FMTs, and the variability of live biotherapeutics.

The extensive variety of recently identified non-canonical conjugated BAs expands the complexity of the BSH-BA pool and reinforces BSHs as key BA-altering enzymes (*16, 17*). BSH-generated deconjugated BAs fuel the production of non-canonical conjugated BAs (**Fig. 4D**). While the mechanism and purpose of this re-conjugation is yet to be elucidated, our observation that non-canonical conjugations can inhibit *C. difficile* spore germination introduces new mechanistic avenues by which BSH activity can be used to restore colonization resistance (**Fig. 4E**). We also demonstrate that BSHs can variably act on these BAs to support a model where BSHs directly shape the non-canonical conjugated BA pool by simultaneously sustaining and curtailing it (**Fig. 4D, S14**). Given the broad impacts BSH can have on the overall BA pool, purposefully augmenting the specificity or activity of BSHs could support their potential use as a precision modality to steer BA metabolism *in vivo*.

BSHs perform the important role of BA deconjugation, and perturbations to this process are implicated in several diseases including CDI. *C. difficile* pathogenesis is exceptionally responsive to alterations in BSH activity, thereby establishing these enzymes as a potential targeted approach to restore colonization resistance. The preferences and efficiency by which BSHs recognize conjugated BAs defines the resulting BA pool, and their ability to variably process the expansive catalogue of non-canonical conjugated BAs is a newly prescribed function of BSHs. In the future, BSHs may be leveraged to rationally manipulate the chemical complexity of the BA pool toward addressing many states of intestinal disease.

## Acknowledgements

This work was performed in part by the Molecular Education, Technology and Research Innovation Center (METRIC) at NC State University, which is supported by the State of North Carolina.

## Author contributions

MHF, MEW, SOF, ECP, MRR, RB, CMT designed the research. MHF, MEW, AKS, SOF, ECG, GA, SP, MP, VVB, MEV, and ESB performed, analyzed, and interpreted experiments. SOF and MP performed bioinformatic analysis of BSHs. MHF and GA performed all BSH specific activity assays and *C. difficile* experiments. MEW, SP, and VVB performed crystallographic work. MD, SKM, and AJ collected patient samples. AKS performed metabolomic analyses. ECG synthesized non-canonical bile acids. MF and MEW wrote the manuscript which all authors edited.

## Data and materials availability

All data associated with this study are available in the main text or the Supplementary Materials. All LC-IMS-MS data are publicly available on Panorama under the Panorama dashboard of the Baker Lab-NCSU within the “0821 BSH Assessment”, “1121 Mouse Germination and Growth Ex-Vivo” and the “1121 Theriot FMT Ex-Vivo Growth” projects.”

## List of Supplementary materials

Materials and Methods

Fig S1 to S17

Table S1 to S3

