## Supplemental Material for "Distinct bile salt hydrolase substrate preferences dictate *C. difficile* pathogenesis"

**This PDF file includes:**

Materials and Methods

Fig S1 to S17

Table S1 to S3

**Materials and Methods**

Phylogenetic analysis of BSH sequences in lactobacilli

The method of BSH sequence analysis in lactobacilli was as previously described (*19*).

Briefly, the whole-genome sequence data for lactobacilli were downloaded from NCBI in July 2021, resulting in a total of 3712 nucleotide files. The data were subsequently built into a local BLAST *Lactobacillus* database. Updated BSH (15 sequences) and penicillin V acylase (PVA) (11 sequences) reference sets were manually created based on previously characterized BSH and PVA proteins (*19*). A BLASTx search was then performed using the BSH reference set as the query and the *Lactobacillus* database as the BLAST database. Search results were filtered using custom code to identify sequences with > 30% identity and at least 100 amino acids in length for downstream analysis.

The BSH and PVA reference sets were separately aligned using MUSCLE (*36*). A Stockholm multisequence alignment (MSA) file, generated using the alignment, was then used to run the Hidden Markov Model (HMM)(*37*) on the results of the BLASTx search to identify likely BSH proteins. Custom code was employed to remove results with E values less than 1e-99 as false positives and to determine whether the amino acid sequence was likely a BSH or PVA based on its E value.

To facilitate the downstream phylogenetic analysis, CD-HIT clustering was used to cluster similar BSH sequences that shared at least 95% identity (*38, 39*). Custom code was employed to extract the consensus sequence from each cluster for the construction of a phylogenetic tree. CD-HIT cluster data were hand-curated to confirm the amino acid sequence of the representative BSH proteins from each of the 84 clusters and to remove redundant sequences.

Chemicals and materials

The following chemicals were used in this study: sodium taurocholate (Cayman Chemical), sodium glycocholate (Sigma Aldrich), sodium taurochenodeoxycholate (Sigma Aldrich), taurine (Sigma Aldrich), glycine (Sigma Aldrich), sodium acetate trihydrate (Fisher), sodium phosphate dibasic dihydrate (Sigma Aldrich), sodium phosphate monobasic dihydrate (Sigma Aldrich), DL-dithiothreitol (MP Biomedicals), trichloroacetic acid (Sigma Aldrich), ninhydrin (Sigma Aldrich), sodium citrate tribasic dihydrate (Sigma Aldrich), glycerol (Fisher), crystal screens MCSG1-4 (Anatrace).

Bacterial strains and growth conditions

BSH-expressing *Escherichia coli* Rosetta (DE3) pLysS were cultured at 37°C shaking overnight in LB broth supplemented with 30 μg mL^-1^ kanamycin (Kan) and 20 μg mL^-1^ chloramphenicol (Cam) or in Terrific Broth (TB) supplemented with the same antibiotics for BSH overexpression. *Lactobacillus* species were cultured statically in MRS media at 37°C in a Coy anerobic chamber (5% H_2_/10% CO_2_/85% N_2_) and *L. gasseri* ATCC 33323 *bsh* mutants were previously described (*8*). *C. difficile* R20291 was statically cultured anaerobically using BHIS broth supplemented with 100 mg/L of L-cysteine. BHIS agar supplemented with 0.1% TCA (TBHIS) was used to start cultures from spore stocks. *C. difficile* enumeration from intestinal contents or stool was performed on CCFA (cefoxitin, cycloserine, and fructose agar) or TCCFA (containing 0.1% TCA). All agar plates were made using 1.5% agar. *C. difficile* growth curves were carried out in BHIS broth in clear flat-bottom plates containing 200 μL of media per well. Growths were performed in a Tecan plate reader within the anaerobic chamber at 37 °C for 24 h.

Recombinant BSH cloning and protein expression

All BSH sequences cloned in this study were amplified from genomic DNA or were codon-optimized in synthesized genes (IDT) using custom oligonucleotide primers (IDT), as reported in **Table S2.** The BSH genes for LaciBSHa, LgasBSHa, LgasBSHb, LjohBSHa, LjohBSHb, LjohBSHc, LcriBSHa, LcriBSHb, and LsalBSH were all amplified from genomic DNA. The genes for LmurBSH, LrogBSH, LplaBSH, LaviBSH, LgigBSH, LreuBSH, LcolBSH, and LingBSH were codon-optimized and synthesized. PCR was performed using Phusion Flash High-Fidelity PCR master mix (Thermo) and products were purified using the QIAquick PCR Purification Kit (Qiagen). Amplicons were cloned into the pETite C-His vecton (Lucigen) were purified using the Monarch Plasmid Miniprep Kit (NEB) and sequenced. Overexpression *E. coli* cultures were grown at 37°C shaking in 1L of TB with Kan and Cam until an OD600 ~ 0.6 was reached. Isopropyl-β-D-thiogalactopyranoside (IPTG) was added to cultures to induce expression and cells were grown at 30°C shaking for 16-20h. Cells were harvested by centrifugation and were stored at -80°C.

Recombinant protein purification

To purify BSHs, frozen *E. coli* cell pellets were resuspended in 50 mL of lysis buffer (50 mM NaPO_4_, 300 mM NaCl, 20 mM imidazole, 10 mM 2-mercapoethanol, protease inhibitor [Roche], DNAse [Sigma], pH 8.0) and were lysed by sonication. Cell debris were pelleted by centrifugation at 25,000 x *g* for 30 min at 4°C. Lysates were run over a gravity column containing 4 mL of fresh HisPur cobalt resin (Thermo Scientific) that was equilibrated in wash buffer (50 mM NaPO_4_, 300 mM NaCl, 20 mM imidazole, pH 8.0). Bound BSHs were washed on the column at a rate of ~1 mL per min with 20 mL of wash buffer and were then eluted with 10 mL of elution buffer, which was wash buffer plus 150 mM imidazole and 10 mM DTT, and flash frozen in liquid N_2_ to prevent oxidation. BSHs were quantified using the Qubit Protein Assay Kit (Invitrogen) and protein purity was assessed using 10%, 14%, or 4-20% SDS-PAGE gels (Thermo Scientific) (**Fig. S2**).

Protein expression and purification for crystallography

The expression plasmid was transformed into BL21-G competent E. coli cells (New England Biolabs) and cultured under kanamycin (25 µg/mL) and chloramphenicol (34 µg/mL) selection. A single colony was selected the next day and grown overnight in 100 mL Terrific Broth media (4.82 g Terrific Broth EZMix powder, 400 µL glycerol, 100 mL DI H2O) at 37°C with shaking at 210 RPM with kanamycin (25 µg/mL) and chloramphenicol (34 µg/mL). The next day, 50 mL of the overnight culture was added to 1.5 L TB media with kanamycin, chloramphenicol, and ~40 µL of Antifoam 204. The culture was then incubated at 37°C with shaking at 210 RPM for expression. Expression was induced when optical density (OD600) reached 0.6-0.8 with 100 µM isopropyl ß-D-1-thiogalactopyranoside (IPTG), and the culture was incubated overnight at 18 °C.

The cells were then pelleted in a Sorvall Instruments RC-3B centrifuge at 4500 xg for 20 min at 4°C. The cell pellet was resuspended in lysis buffer (30 mL BSH Buffer A: 50 mM sodium phosphate, 300 mM sodium chloride, 10 mM imidazole, pH 8.0 with one cOmplete Mini EDTA-free protease inhibitor cocktail tablet (Roche), lysozyme, 50 µg/mL DNAse, and 10 mM 2-mercaptoethanol). Cells were then lysed by sonication on ice using a Fischer Scientific Sonic dismembrator model 500 with 0.5 s pulses for 1.5 min three times at 40%, 50%, and 60% intensity. The cell suspension was pelleted by centrifugation in a Beckman Coulter J2-HC centrifuge at 17,000 xg for 50 min at 4 °C. The supernatant was syringe-filtered using a 0.22 µm filter.

The filtrate was flowed over a HisTrap HP 5 mL (Cytiva) column using the Aktaexpress FPLC (Amersham Bioscience) and washed with BSH buffer A. The bound protein was eluted in a stepwise fashion with 100% BSH Buffer B (50 mM sodium phosphate, 300 mM sodium chloride, 300 mM imidazole, pH 8.0). Fractions containing the protein of interest were combined and flowed over a HiLoad 16/60 Superdex 200 gel filtration column (GE Life Sciences). Samples were eluted in S200 sizing buffer (20 mM HEPES, 50 mM NaCl, pH 7.4). Fractions containing the protein of interest were analyzed for purity using SDS-PAGE, and those with >95% purity were combined and concentrated using 30 kDa cutoff molecular weight centrifuge concentrators (EMD Millipore) to ~10 mg/mL. The final protein concentration was determined using a ND-1000 spectrophotometer. DL-dithiothreitol (DTT) was added to the concentrated fractions at a final concentration of 10 mM, and samples were snap-frozen in liquid nitrogen and stored at -80°C.

BSH specific activity assays

Activity assays were performed as previously described (*8*). Briefly, the assay reacted 10-25 nM BSH with 9 mM conventional BAs for 5 min, or 100 nM BSH with 4.5 mM non-canonical BAs/penV, for 1 h in 50 μL volumes. Reactions were carried out in 0.1 mM Na phosphate, 10 mM DTT, pH 6.0 and stopped with 50 μL of trichloroacetic acid. To determine the quantity of amino acid or 6-APA released, the colorimetric ninhydrin reaction was carried out. 25 μL of the quenched BSH reaction was added to 475 μL of ninhydrin buffer (0.3 mL glycerol, 0.175 mL 0.5 M Na Citrate pH 5.5, 0.25% ninhydrin reagent) and boiled for 14 min. A standard curve of the respective conjugated amino acid or 6-APA was prepared for each assay. Absorbance was measured at 570 nm in clear flat-bottom plates in a Tecan Infinite F200 Pro plate reader. Specific activity is reported as μmol amino acid released / s / μmol BSH.

Protein crystallography

Crystals were generated at 20ºC using sitting drop vapor diffusion for all specimen obtained except those of LgasBSHa grown with taurine and CDCA, which employed hanging drop vapor diffusion. Sitting drop trays were set up using the Douglas Instruments Oryx4 instrument and Hampton Research 3-well midi crystallization plates (Swissci). Hanging drops were manually plated using EasyXtal 15-well Tool trays (Qiagen). Specific crystallization conditions follow. LaciBSHa: Crystallant composed of 0.15 M DL-malic acid, pH 7.0, with 20% (w/v) PEG 3350. Crystals formed in a 2:1 protein (11.57 mg/mL) to crystallant ratio. LgasBSHa with taurine: Crystallant composed of 0.2M potassium sulfate, 20% (w/v) PEG 3350. Crystals formed in a 1:2 protein (9.5 mg/mL) to crystallant ratio. LgasBSHa with taurine and CDCA: A condition containing 18% (w/v) PEG 3350, 0.2M ammonium chloride pH 6.3 in a hanging drop tray was streak seeded with crystals grown using the sitting drop method in 20% (w/v) PEG 3350, 0.2M ammonium chloride pH 6.3. Crystals were grown in a 1:2 protein (9.55 mg/mL) to crystallant ratio. The resultant crystals were soaked in 1:1 ratio of crystallant:100 mM taurochenodeoxycholic acid for 24h before looping. LgasBSHb: Crystallant composed of 0.2 M magnesium chloride, 0.1 M sodium cacodyolate:HCl, pH 6.5, 20% (w/v) PEG 1000. Crystals formed in a 1:2 protein (10.75 mg/mL) to crystallant ratio. LjohBSHc: Crystallant composed of 0.1 M sodium citrate:HCl, pH 5.6, 10% (w/v) PEG 4000, 10% (v/v) isopropanol. Crystals formed in a 2:1 protein (10.34 mg/mL) to crystallant ratio. LingBSH: Crystallant composed of 0.2 M calcium acetate hydrate, 0.1M Tris:HCl pH 7.0, 20% (w/v) PEG 3000. Crystals formed in a 1:2 protein (13.8 mg/mL) to crystallant ratio. LreuBSH: Crystallant composed of 0.1 M Bis-Tris Propane:HCl, pH 7.0, 1.5 M ammonium sulfate. Crystals formed in a 2:1 protein (13.3 mg/mL) to crystallant ratio.

Crystals were cryo-protected in the conditions described above with 20% glycerol before looping and flash cooling in liquid nitrogen. Diffraction data were collected at 100K at either APS 23ID-D (LgasBSHa with taurine and CDCA, LingBSH), APS 23ID-B (LaciBSHa, LgasBSHa with taurine, LgasBSHb, LjohBSHc), or ALS 5.0.2 (LreuBSH). Data were processed and scaled with XDS (*40*), and Phenix (version 1.17.1-3660) was employed to perform molecular replacement with the model PDB 2HEZ. Output models from Phaser were improved with Autobuild (*41*) and structures were refined using phenix.refine (*41*) with iterative cycles of manual adjustment using Coot (version 0.9.4.1) (*42*). Final coordinates were deposited into the RCSB Protein Data Bank under the codes 7SVE [10.2210/pdb7sve/pdb], 7SVF [10.2210/pdb7svf/pdb], 7SVG [10.2210/pdb7svg/pdb], 7SVH [10.2210/pdb7svh/pdb], 7SVI [10.2210/pdb7svi/pdb], 7SVJ [10.2210/pdb7svj/pdb], and 7SVK [10.2210/pdb7svk/pdb].

Glycine modelling

Glycine was modelled into the active site of the unliganded structure of LgasBSHb by superimposing this enzyme’s active site with that of the structure of LgasBSHa in complex with both taurine and CDCA, with glycine manually positioned in place of taurine using PyMOL (version 2.3.2).

BSH catalytic efficiency

Assays were performed by adding 10 µL enzyme (10-100 nM final, depending on enzyme) to 30 µL of substrate at 4 concentrations (taurocholic acid or glycocholic acid, 500-1500 µM final concentration) and 10 µL assay buffer (50 mM sodium acetate pH 5.0, 10 mM dithiothreitol for LgasBSHb, 50 mM sodium phosphate pH 6.5, 10 mM dithiothreitol for LgasBSHa) for a total volume of 50 µL. Reactions were incubated at 37ºC and quenched at five time points with 50 µL 15% (w/v) trichloroacetic acid. Reactions were centrifuged at 4000g for 2 minutes. To quantitate the amount of liberated amino acid present, 10 µL quenched reaction was added to 190 µL ninhydrin reaction (62.5 mL of 1% ninhydrin in 0.5 M sodium citrate [pH 5.5], 150 mL of glycerol, 25 mL of 0.5 M sodium citrate [pH 5.5]) in a 96-well PCR plate. A standard curve of glycine or taurine was also created using the ninhydrin reaction. Using a thermocycler, the mixtures were heated to 90ºC for 14 minutes and then cooled at 25ºC for 15 minutes. 150 µL ninhydrin mixture was then added to a 96 well flat black plate (Grenier 655906) and the absorbance was measured at 570 nm using a Clariostar Plus plate reader (BMG Labtech). The standard curve was used to quantitate the amount of free amino acid liberated by the reaction. Reaction curves over time were fit using linear regression, and the resultant initial velocities were used to determine k_cat_/K_M_ by plotting velocity against substrate concentration. Catalytic efficiencies are the average of three biological replicates ± SD.

Spore preparation

*C. difficile* spores were prepare as previously described (*31, 43*). Briefly, *C. difficile* was grown at 37°C anaerobically for 1 week in Clospore media (*44*). Spores were harvested by centrifugation and washed with water, heat treated for 20 min ad 65°C, and were stored at 4°C. Spores were plated on BHIS and TBHIS to make sure no viable cells were present.

Spore germination and outgrowth assays

Spore germination experiments were modified from Carlson et al. and Theriot et al. (*43, 45*). Conventional BAs were dissolved in water and non-canonical BAs were dissolved in methanol. For the inhibition of germination experiments, the germinant TCA was used across conditions at 0.1% (1.9 mM). CDCA was used at 0.04% (0.965 mM) as a positive control for germination inhibition and all other BAs were tested at the same concentration (0.965 mM). Germination reactions were carried out anaerobically in 100 μL volumes in PBS for 30 min. Reactions were subsequently serially diluted and plated on TBHIS and BHIS agar. Percent germination for each BA was calculated as [100 x (CFUs on BHIS / CFUs on TBHIS)] and was normalized to germination with TCA alone. For the germination reactions with non-canonical BAs, all BAs were used at 1.9 mM and percent germination was calculated and normalized as stated above.

Minimum inhibitory concentrations

BA tolerance measured by minimum inhibitory concentration was adapted from the bile tolerance assay by Jacobsen et al. (87). Overnight *C. difficile* cultures (~10^8^ CFUs/mL) were inoculated 1% into BHIS containing a range of BA concentrations. Cultures were anaerobically incubated for 24 h at 37 °C. Following incubation, cultures were serially diluted in PBS and plated on BHIS agar to determine if the concentration of BA tested inhibited growth relative to the starting inoculum.

Propidium iodide (PI) staining

Exponentially growing *C. difficile* cultures were captured and washed 3 times with PBS before being back diluted to a final OD600 = 0.1 into PBS containing BAs at 0.25 x the MIC for 30 min. Bacteria were exposed to 150 μM SDS as positive controls for detergent induced membrane damage and PI staining. Following ΒΑ exposure, bacteria were stained for 30 min at 37 °C with 20 μg/mL PI using slight modifications to a previous method (*46, 47*). Stained bacteria were diluted 1:10 in PBS and PI fluorescence was measured from flat-bottom clear plates (excitation: 540 nm; emission: 610 nm) in 100 μl volumes in a Tecan Infinite F200 Pro plate reader and background fluorescence was subtracted.

Pre-FMT sample collection

All consenting patients undergoing FMT for rCDI at the University of North Carolina from January to December 2017 in a prospective registry were enrolled for fecal collection. rCDI was defined by a patient having at least the third episode of CDI. There were no exclusion criteria for participation in the registry specifically, though subjects were by definition undergoing FMT under the care of a physician who judged the benefits to outweigh the risks. Patient stool samples from 2 weeks prior to FMT (pre-FMT) were collected. The study was approved by the UNC IRB (#16-2283).

Animals and housing

Animal experiments were conducted in the Laboratory Animal Facilities located on the North Carolina State University (NCSU) College of Veterinary Medicine (CVM) campus. The animal facilities are equipped with a full-time animal care staff coordinated by the Laboratory Animal Resources division at NCSU. The NCSU CVM is accredited by the Association for the Assessment and Accreditation of Laboratory Animal Care International. The Institutional Animal Care and Use Committee at the NCSU CVM approved this study. Trained animal handlers in the facility fed and assessed the status of animals several times per day.

Mouse sample collection

6-8 week old male C57BL/6 mice (n = 16 total) were given 0.5 mg/mL cefoperazone in their drinking water for 5 days to make them susceptible to *C. difficile* infection (*48*). Subsequently, the mice were then given normal water (Gibco) for 2 days, after which groups of mice (n = 3-4) were sacrificed at days 0, 2, 4, and 7. Small intestinal and cecal contents from each mouse were collected and samples were flash frozen in liquid N_2_.

*Ex vivo* growth and spore germination assay

To measure BSH-dependent inhibition of *C. difficile* *ex vivo* in pre-FMT stool and mouse intestinal contents, frozen material was thawed and diluted 1:3 in PBS. For growth experiments, pre-FMT stool samples were treated with PBS, the taurine-preferring BSH cocktail T-BSH (LgasBSHa, LjohBSHc, and LsalBSH), the glycine-preferring BSH cocktail G-BSH (LjohBSHa, LgasBSHb, and LingBSH), or the combined broad acting G&T-BSH cocktail for 30 min at 37 °C. Mouse contents were treated with PBS or the G&T-BSH cocktail. All BSHs had a final concentration of 0.1 μM each. Immediately afterwards, exponentially growing *C. difficile* was inoculated into samples to a final concentration of 10^5^ CFUs/mL. *C. difficile* was cultured anaerobically at 37 °C and enumerated on TCCFA at 8 and 24 h.

To measure BSH-dependent inhibition of germination in mouse small intestinal contents, 1:3 diluted samples were treated with PBS or the G&T-BSH cocktail for 8 h at 37 °C. 10^6^ *C. difficile* spores/mL were then inoculated in the small intestinal contents for 30 min and samples were promptly diluted and enumerated on CCFCA and TCCFA. % germination was calculated as [100% x (CFUs on CCFA / CFUs on TCCFA)].

Non-canonical bile acid synthesis

All non-canonical conjugated BAs were purified according to literature precedent and characterization data is consistent with reported data (*16, 18, 49, 50*). Synthesis of the conjugated bile acids was adapted from a method published by Ezawa *et al (51)*.  Bile acid (0.25 mmol, 1 equiv) was dissolved in anhydrous THF (4.9 mL, 0.05M) and cooled to 0°C with stirring. Ethyl chloroformate (0.3 mmol, 1.2 equiv) was added followed by Et_3_N (0.3 mmol, 1.2 equiv) and the reaction was stirred for 1.5h at 0°C. After conversion of starting material by TLC, a cold solution of amino acid (0.375 mmol, 1.5 equiv) and either NaHCO_3_ or NaOH (0.375 equiv, 1.5 equiv) in H_2_O (4.9 mL, 0.05M) was added in one portion.  Then, the reaction was stirred for 2 h, allowing it to gradually warm to rt. After this time, THF was removed *in vacuo*, then 2M HCl was added to acidify the reaction mixture to pH<2, producing a white precipitate.  The mixture was extracted in ethyl acetate (3 ✕ 20 mL), then the combined organic layers were washed with brine (50 mL), dried over Na_2_SO_4_, and concentrated *in vacuo*.  The crude residue was purified over silica gel using methanol and dichloromethane with 1% acetic acid.

Organic solutions were concentrated under reduced pressure on a Büchi rotary evaporator using a water bath. Chromatographic purification of products was accomplished by flash chromatography on Silicycle F60 silica gel. All reactions were carried out in well ventilated fume hoods. Thin-layer chromatography (TLC) was performed on Silicycle 250 μm silica gel plates. Visualization of the developed chromatogram was performed by irradiation with 254 nm UV light or treatment with a solution of ceric ammonium molybdate stain followed by heating. Yields refer to purified compounds unless otherwise noted.

^1^H and ^13^C NMR spectra were recorded on a Bruker 600 (600 and 151 MHz for ^1^H and ^13^C, respectively) instrument, and are internally referenced to residual protiosolvent signals of CD_3_OD at δ 3.31 and 49.00 and (CD_3_)_2_SO at 2.50 and 39.52. Data for ^1^H NMR are reported as follows: chemical shift (δ ppm), integration, multiplicity (s = singlet, br s = broad singlet, d = doublet, t = triplet, q = quartet, m = multiplet), and coupling constant (Hz). Data for ^13^C NMR are reported in terms of chemical shift and no special nomenclature is used for equivalent carbons.

Glu-CDCA**,** Ile-CA, Leu-CA, Phe-CA, Trp-CDCA and Tyr-CA: Purified according to literature precedent and characterization data is consistent with reported data (*16, 18*).

Ala-CA: NaHCO_3_ was used as the inorganic base. Product was purified using 3-10% CH_3_OH in CH_2_Cl_2_ with 1% acetic acid to obtain a 92% yield as a white amorphous solid. ^1^H NMR (600 MHz, CD_3_OD) δ 4.40 – 4.29 (m, 1H), 3.96 (t, *J* = 3.1 Hz, 1H), 3.80 (q, *J* = 3.1 Hz, 1H), 3.41 – 3.36 (m, 1H), 2.33 – 2.21 (m, 3H), 2.20 – 2.12 (m, 1H), 2.00 – 1.93 (m, 2H), 1.92 – 1.77 (m, 4H), 1.77 – 1.70 (m, 1H), 1.69 – 1.63 (m, 1H), 1.63 – 1.50 (m, 5H), 1.47 – 1.40 (m, 2H), 1.38 (d, *J* = 6.9 Hz, 3H), 1.36 – 1.24 (m, 3H), 1.11 (qd, *J* = 11.7, 5.5 Hz, 1H), 1.03 (d, *J* = 6.6 Hz, 3H), 0.98 (td, *J* = 14.2, 3.3 Hz, 1H), 0.91 (s, 3H), 0.71 (s, 3H). ^13^C NMR (151 MHz, MeOD) δ 176.46, 74.01, 72.83, 69.02, 49.85, 48.04, 47.46, 43.14, 42.93, 40.97, 40.40, 36.83, 36.47, 35.86, 35.82, 33.85, 33.09, 31.13, 29.52, 28.65, 27.81, 24.21, 23.18, 17.80, 17.76, 13.01. HRMS(ESI) exact mass calculated for [M+H]^+^ (C_27_H_46_NO_6_) requires m/z 480.3320, found 480.3320 with a difference of 0.00 ppm.

His-CDCA: NaOH was used as the inorganic base. Product was purified using 3-10% CH_3_OH in CH_2_Cl_2_ with 1% acetic acid to obtain a 41% yield as a white amorphous solid. ^1^H NMR (600 MHz, CD_3_OD) δ 8.52 (s, 1H), 8.24 (s, 1H, NH), 7.20 (s, 1H), 4.58 (dd, *J* = 8.0, 5.1 Hz, 1H), 3.79 (q, *J* = 3.0 Hz, 1H), 3.38 (tt, *J* = 11.7, 4.6 Hz, 1H), 3.24 (dd, *J* = 15.1, 5.2 Hz, 1H), 3.05 (dd, *J* = 15.1, 8.0 Hz, 1H), 2.34 – 2.21 (m, 2H), 2.20 – 2.08 (m, 1H), 2.03 – 1.93 (m, 2H), 1.93 – 1.81 (m, 3H), 1.78 – 1.69 (m, 2H), 1.69 – 1.57 (m, 2H), 1.57 – 1.40 (m, 5H), 1.40 – 1.22 (m, 5H), 1.21 – 1.06 (m, 3H), 1.02 – 0.96 (m, 1H), 0.95 (d, *J* = 6.5 Hz, 3H), 0.93 (s, 3H), 0.68 (s, 3H). ^13^C NMR (151 MHz, CD_3_OD) δ 176.35, 175.72, 132.60, 118.14, 72.84, 69.06, 57.31, 54.33, 51.56, 43.67, 43.15, 41.06, 40.74, 40.46, 36.96, 36.54, 36.21, 35.91, 34.06, 34.00, 33.10, 31.35, 29.27, 24.63, 23.39, 21.78, 18.92, 12.18. HRMS (ESI) exact mass calculated for [M+H]+ (C_30_H_48_N_3_O_5_) requires m/z 530.3589, found 530.3591 with a difference of 0.38 ppm.

Ser-CA: NaHCO_3_ was used as the inorganic base. Product was purified using 6-12% CH_3_OH in CH_2_Cl_2_ with 1% acetic acid to obtain a 74% yield as an off-white amorphous solid. ^1^H NMR (600 MHz, CD_3_OD) δ 4.33 (s, 1H), 3.95 (t, *J* = 3.1 Hz, 1H), 3.88 – 3.75 (m, 3H), 3.41-3.36 (m, 1H), 2.40 – 2.33 (m, 1H), 2.31 – 2.16 (m, 3H), 2.03 – 1.98 (m, 1H), 1.95 – 1.71 (m, 7H), 1.69 – 1.63 (m, 1H), 1.63 – 1.51 (m, 5H), 1.47 – 1.35 (m, 4H), 1.34 – 1.26 (m, 1H), 1.11 (qd, *J* = 11.7, 5.5 Hz, 1H), 1.04 (d, *J* = 6.4 Hz, 3H), 0.98 (td, *J* = 14.1, 3.3 Hz, 1H), 0.91 (s, 3H), 0.71 (s, 3H). ^13^C NMR (151 MHz, CD_3_OD) δ 176.47, 74.03, 72.88, 69.04, 63.77, 49.85, 48.04, 47.50, 43.20, 42.99, 41.02, 40.46, 37.00, 36.49, 35.90, 35.86, 34.17, 33.04, 31.18, 29.58, 28.72, 27.88, 24.24, 23.17, 17.80, 13.01. HRMS (ESI) exact mass calculated for [M+H]^+^ (C_27_H_46_NO_7_) requires m/z 496.3269, found 496.3267 with a difference of 0.40 ppm.

Trp-CA: NaOH was used as the inorganic base. Product was purified using 6-12% CH_3_OH in CH_2_Cl_2_ with 1% acetic acid to obtain 49% yield as an off-white amorphous solid. ^1^H NMR (600 MHz, CD_3_OD) δ 7.58 (d, *J* = 7.9 Hz, 1H), 7.34 (d, *J* = 8.1 Hz, 1H), 7.14 – 7.07 (m, 2H), 7.02 (t, *J* = 7.4 Hz, 1H), 4.75 (dd, *J* = 8.4, 4.9 Hz, 1H), 3.93 (t, *J* = 3.1 Hz, 1H), 3.81 (q, *J* = 3.1 Hz, 1H), 3.42 – 3.35 (m, 2H), 3.17 (dd, *J* = 14.7, 8.4 Hz, 1H), 2.34 – 2.20 (m, 3H), 2.14 – 2.06 (m, 1H), 2.03 – 1.93 (m, 2H), 1.86 – 1.76 (m, 3H), 1.76 – 1.65 (m, 3H), 1.64 – 1.50 (m, 6H), 1.49 – 1.41 (m, 1H), 1.41 – 1.29 (m, 2H), 1.28 – 1.15 (m, 2H), 1.13 – 1.05 (m, 1H), 1.05 – 1.00 (m, 1H), 0.98 (d, *J* = 6.7 Hz, 3H), 0.92 (s, 3H), 0.67 (s, 3H). ^13^C NMR (151 MHz, CD_3_OD) δ 176.70, 175.49, 137.97, 128.83, 124.30, 122.35, 119.78, 119.21, 112.27, 111.07, 74.04, 72.84, 69.08, 54.62, 47.94, 47.40, 43.10, 42.90, 40.91, 40.40, 36.71, 36.43, 35.84, 35.80, 33.79, 32.96, 31.12, 29.47, 28.54, 28.45, 27.80, 24.19, 23.14, 17.70, 12.96. HRMS (ESI) exact mass calculated for [M+H]^+^ (C_35_H_51_N_2_O_6_) requires m/z 595.3742, found 595.3737 with a difference of 0.84 ppm.

Non-canonical AA-BA pooled activity assays

BSH activity with the pooled AA-BAs was carried out using similar reactions condition as described above in the BSH activity assays with purified BAs. 100 nM BSH was reacted with (2.5 mg/mL) AA-CA, AA-CDCA, AA-DCA, or AA-βMCA, for 1 h in 50 μL volumes in triplicate. 10 μL samples were taken at 5 min, 30 min, and 1 h and immediately quenched by diluting them into 90 μL of methanol and stored at -80 °C. Reactions containing no BSH were used as a negative control. % deconjugation was calculated at each time as follows:

$\boldsymbol{\%}Deconjugation=100\% \times\frac{(BA peak area at 0min) - \left( BA peak area at 5/30/60 min \right)}{BA peak area at 0 min}$

*L. gasseri* whole cell activity assays were carried out in a similar manner with identical reaction conditions. *L. gasseri* wild-type, Δ*bshA*, Δ*bshB*, and Δ*bshAB* cultures were grown to mid-log, washed 3x in PBS, and were diluted to a final concentration of OD600 = 0.1 in reactions containing AA-CDCA. After 1 h min, 50 μL samples was taken and cells were quickly pelleted before 10 μL of supernatant was diluted into 90 μL of methanol and stored at -80 °C. To calculate % deconjugation in the wild-type, Δ*bshA*, or Δ*bshB* conditions, each conjugated BA was normalized to its abundance in the the Δ*bshAB* condition to control for the absorption/adsorption of BAs to the *L. gasseri* cell as follows:

$\boldsymbol{\%}Deconjugation=100\% \times\frac{(BA peak area in \Delta bshAB\mathrm{condition}) - \left( BA peak area in WT/\Delta bshA/\Delta bshB\mathrm{condition} \right)}{BA peak area in \Delta bshAB \mathrm{condition}}$

Bile acid metabolomics

Metabolomic analysis of untreated pre-FMT samples were performed by Metabolon, Inc., in the same manner as was described in our previous study (*52*). Briefly, individual samples were subjected to methanol extraction and then split into aliquots for analysis by ultra-high-performance liquid chromatography-mass spectrometry (UHPLC/MS). The global biochemical profiling analysis comprised four unique arms consisting of reverse-phase chroma- tography positive-ionization methods optimized for hydrophilic compounds (LC/MS Pos Polar) and hydrophobic compounds (LC/MS Pos Lipid) and reverse-phase chromatography performed under negative-ionization conditions (LC/MS Neg) as well as a hydrophilic interaction liquid chromatography (HILIC) method coupled to negative ionization (LC/MS Polar) . All the methods alternated between full-scan MS and data-dependent MS*n* scans. The scan ranges differed slightly between methods but generally covered 70 to 1,000 *m*/*z*.

Metabolites were identified by automated comparison of the ion features in the experimental samples to a reference library of chemical standard entries that included retention time, molecular weight (*m*/*z*), preferred adducts, and in-source fragments as well as associated MS spectra and were curated by visual inspection for quality control using software developed at Metabolon. Identification of known chemical entities was based on comparisons to metabolomic library entries of purified standards (*53*).

For meteabolomic analysis of treated pre-FMT samples and murine intestinal contents, thawed *ex vivo* samples were homogenized using a genie disrupter (Scientific Industries) for 15 min after vortex mixing for 30 min. Samples were then centrifuged at 15,000 rpm and 4 °C for 10 min and the supernatant was removed and used for analysis. 10 μL of the extracted sample was added onto filter spots suspended in the wells of a 96-well filter plate (PALL AcroPrep, PTFE 0.2 μm) fixed on top of a deep-well plate (Waters QuanRecovery) and extracted with 100 μL methanol by shaking at 600 rpm for 20 min. Elution of the methanol extracts was performed using a positive-pressure manifold (Waters) into the lower receiving deep-well plate, which was then detached from the upper filter plate.

After adding 50 μL MS grade water to the extracts and shaking briefly (600 rpm, 5 min), samples were injected (5 uL) directly from the deep well plate. Samples from the non-canonical AA-BA pooled activity assays were injected (5 uL) without dilution or extraction. LC-IMS-MS analyses were performed using an Agilent 1290 Infinity UPLC system (Agilent Technologies, Santa Clara, CA, USA) coupled with an Agilent 6560 IM-QTOF MS instrument (Agilent Technologies, Santa Clara, CA, USA). Chromatographic separation was achieved using a Restek Raptor C18 column (1.7 μm, 2.1 x 50 mm) heated to a temperature of 60°C. Mobile phase A was comprised of 5 mM ammonium acetate, while mobile phase B was 1:1 methanol/acetonitrile. The LC initially started at 0.5 mL/min and 15% B with isocratic elution for 2 minutes followed by a stepped gradient going to 80% B over the next 7.7 min (15-35% B over 2 min, 35-40% B over 2 min, 40-50% B over 1.5 min, 50-55% B over 1.1 min, 55-80% B over 1.1 min). The flow rate and mobile phase composition were then increased to 0.8 mL/min and 85% B for an additional 0.8 minutes followed by re-equilibration at initial conditions for 2 min, resulting in a total run time of 12.5 minutes. Samples were analyzed with negative ion mode (capillary voltage 4000V, nebulizer gas pressure 40 psi, ion source temperature 325°C, dry gas flow 10 L/min) and data were collected from 50-1700 m/z with an IMS drift potential of 17.2 V/cm, frame rate of 0.09 Frames/s, IMS transient rate of 16 IMS transients/frame, maximum IMS drift time of 60 ms and TOF transient rate of 600 transients/IMS transient. LC-IMS-MS spectra were acquired using MassHunter Acquisition Software and raw files (.d) were uploaded to Skyline^1^ for peak picking and molecular annotation using a library built from the synthesized bile acid mixtures and standard mixtures. Peak areas per compound per sample were exported to excel and utilized for statistical analysis in GraphPad Prism. All LC-IMS-MS data are publicly available on Panorama under the Panorama dashboard of the Baker Lab-NCSU within the “0821 BSH Assessment”, “1121 Mouse Germination and Growth Ex-Vivo” and the “1121 Theriot FMT Ex-Vivo Growth” projects.

Statistical analysis

All statistical analysis was performed in GraphPad Prism 8 or 9. Catalytic efficiency assays were analyzed as the average of n = 3 experiments with a two-way ANOVA with Tukey’s multiple comparisons test. Comparisons were made separately between the wild type and mutant versions of LgasBSHa and LgasBSHb. Inhibition of *C. difficile* germination and membrane integrity assays were analyzed from an n = 3 experiments with a one-tailed Welch’s *t* test. Statistical comparisons were only made between conditions containing related BAs (i.e. TCA, GCA, and CA). *C. difficile* growth and bile acid metabolomics in pre-FMT samples were analyzed from an n=3 experiments using a one-way repeated measures ANOVA with Sidak’s correction for multiple comparisons. In the cases where CFU or bile acid metabolomic data was generated from intestinal contents that were split and treated with PBS or a BSH cocktail, one-tailed ratio paired *t* tests or were used to analyze findings. All graphed bars represent mean ± standard deviation. Asterisks indicate significant differences (*p < 0.05, **p < 0.01, ***p < 0.001, ****p < 0.0001).

**Supplemental figure legends:**

**Figure S1. Bile acid structures and abbreviations.**

**Figure S2. SDS-PAGE of purified *Lactobacillus* BSHs.** Protein ladder sizes noted in kDa to the left of gels.

**Figure S3. Structural and sequence comparisons of *Lactobacillus* BSHs.**

(A) Full tetrameric structures of the crystal structures presented here and colored as indicated in **Fig. 2B** (B) Root-mean square deviation (RMSD) in Å across equivalent Cα positions for the crystal structures presented, as well as % sequence identity values.

**Figure S4. BSH multiple sequence alignment.**

Multiple sequence alignment of the Lactobacilli BSH proteins examined here. The three-residue selectivity region is highlighted in yellow (glycine-preferring enzymes) or blue (taurine-preferring enzymes). Created using the ClustalOmega Multiple Sequence Alignment tool.

**Figure S5. Bile acid deconjugations differentially disrupt *C. difficile* membrane integrity.** Propidium iodide staining of exponentially growing *C. difficile* after a 30 min exposure to BAs (n = 3). BA concentrations were 0.25x their MIC. The detergent SDS (150 μM) was used as a positive control for detergent-induced membrane damage. Asterisks indicate significant differences (**p* < 0.05, ***p* < 0.01) between BAs by one-tailed Welch’s *t* test. Comparisons were only made between BAs that share the same BA sterol core. Bars represent mean ± standard deviation.

**Figure S6. Bile acid metabolomics of pre-FMT stool**. Bile acids identified in pre-FMT recipient’s stool prior to BSH treatment.

**Figure S7. *C. difficile* growth in pre-FMT stool**. Growth was measured at 8 and 24 h from an n = 3 replicates. Targeted metabolomics showing the deconjugated BAs CA and CDCA from the samples from. Bars represent mean ± standard deviation. Asterisks indicate significant differences (**p* < 0.05, ***p* < 0.01, ****p* < 0.001, *****p* < 0.0001) between CFUs by one-way repeated measures ANOVA with Sidak’s correction for multiple comparisons.

**Figure S8. *C. difficile* growth with BSH cocktail.** *C. difficile* growth is not impacted by the amino acids (4.5 mM each) or the BSH cocktail.

**Figure S9. Bile acid metabolomics of BSH-treated pre-FMT stool.** Targeted metabolomics showing all detected BAs at 8 h in pre-FMT stool samples. Bars represent mean ± standard deviation. Asterisks indicate significant differences (**p* < 0.05, ***p* < 0.01, ****p* < 0.001, *****p* < 0.0001) from PBS by one-way repeated measures ANOVA with Sidak’s correction for multiple comparisons.

**Figure S10. Growth of *C. difficile* at 24h in murine intestinal content.** Contents (n = 3-4 mice) were treated with PBS or a BSH cocktail and inoculated simultaneously. Asterisks indicate significant differences (**p* < 0.05, ***p* < 0.01, ****p* < 0.001) between BAs by ratio paired one-tailed *t* test. Bars represent mean ± standard deviation.

**Figure S11. Bile acid metabolomics of *C. difficile* spore germination and growth in mouse intestinal content.** Asterisks indicate significant differences (**p* < 0.05, ***p* < 0.01, ****p* < 0.001, *****p* < 0.0001) between treatments by ratio paired one-tailed *t* test. Bars represent mean ± standard deviation from n = 4 samples.

**Figure S12. *C. difficile* growth and spore germination with non-canonical conjugated BAs.** (A) Growth was carried out in the presence of 2 mM BAs in triplicate. Trp-CDCA inhibited growth at this concentration and its MIC was determined to be 0.5 mM. (B) Germination of *C. difficile* was carried out using 2 mM BAs. Germination inhibition with the germinant TCA (2 mM) was carried out with 0.75 mM BA. Bars represent mean ± standard deviation. Asterisks indicate significant differences (****p* < 0.001, *****p* < 0.0001) from TCA only by one-way repeated measures ANOVA with Sidak’s correction for multiple comparisons.

**Figure S13. BSH activity with purified non-canonical bile acids.** Heatmaps of BSH specific activity using purified non-canonical conjugated BAs. Values represent mean activity (n = 3). BSHs were arranged into a phylogeny based on their amino acid sequence. The ability to process penV was included as a potential BSH substrate.

**Figure S14.** **Broadly surveying BSH processing of non-canonical conjugated BAs.**

(A) Schematic of BSH reactions with pooled non-canonical conjugated BAs. Individual BSHs were reacted with pooled BA over an hour. Reactions were halted and BA abundance was quantified using LC-IMS-MS targeted BA metabolomics to measure deconjugation over time. (B) A heatmap of BSH activity across all non-canonical BA pools. Values represent the mean from n = 3 replicates of % deconjugation after 5 minutes, which was calculated by measuring the abundance of an individual BA at 0m and 5m to quantify the proportion of substrate deconjugated. (C) A heatmap of *L. gasseri* BSH activity after 1h.

**Figure S15. LingBSH deconjugation kinetics of non-canonical conjugated bile acids.** LingBSH-catalyzed deconjugation of non-canonical conjugated BAs at 5, 30, and 60 min. Lines are plotted from mean from a n = 3 replicates.

**Figure S16. LingBSH activity compared to non-canonical conjugated amino acid size.** Initial activity of LingBSH with AA-BA mixtures. All dots represent the mean deconjugation from a n = 3 replicates ± standard deviation. individual non-canonical BAs. The MW of Gly and Tau are indicated on the x-axis for reference.

**Figure S17. ^1^H and ^1^3C NMR spectra of pure synthesized conjugated bile acids.**

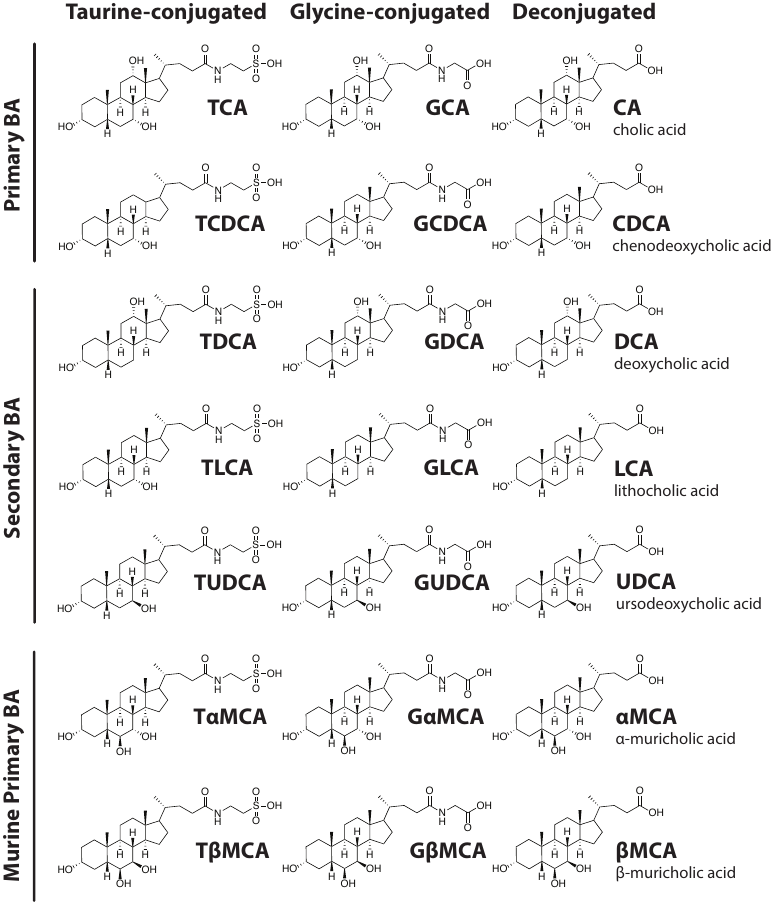

**Figure S1**

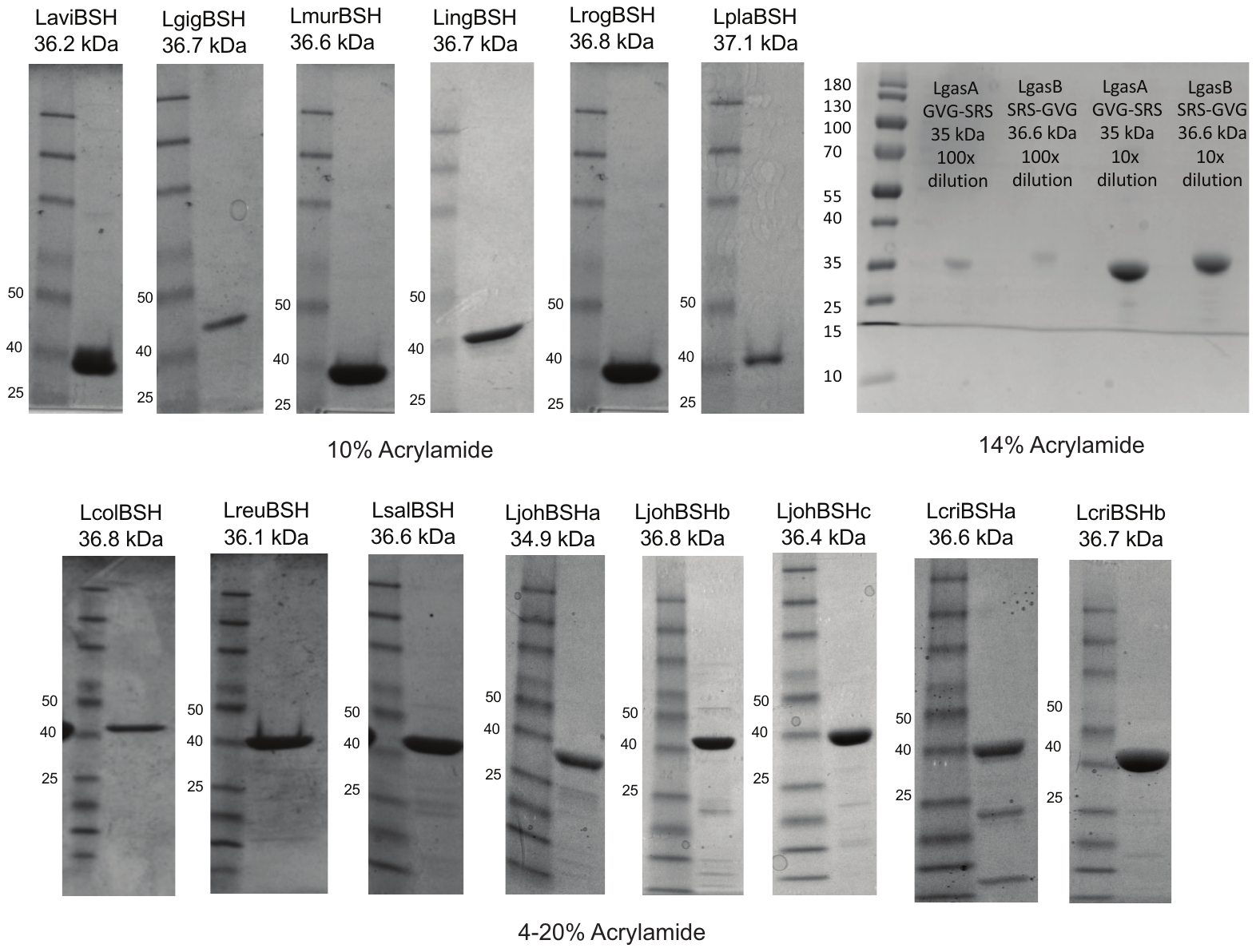

**Figure S2**

**Figure S3**

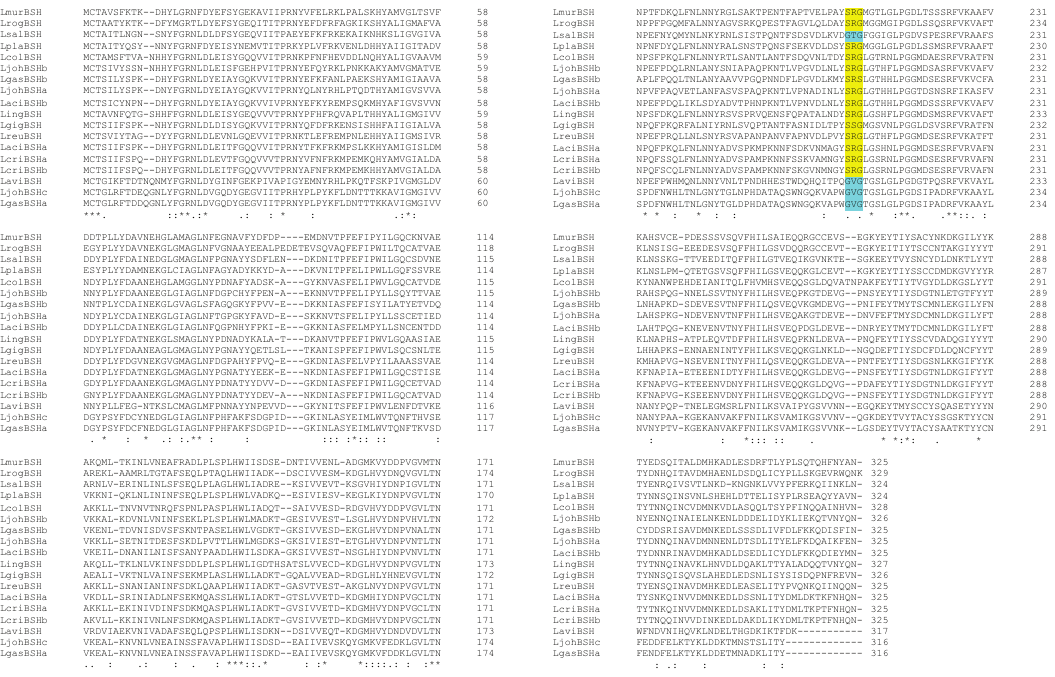

**Figure S4**

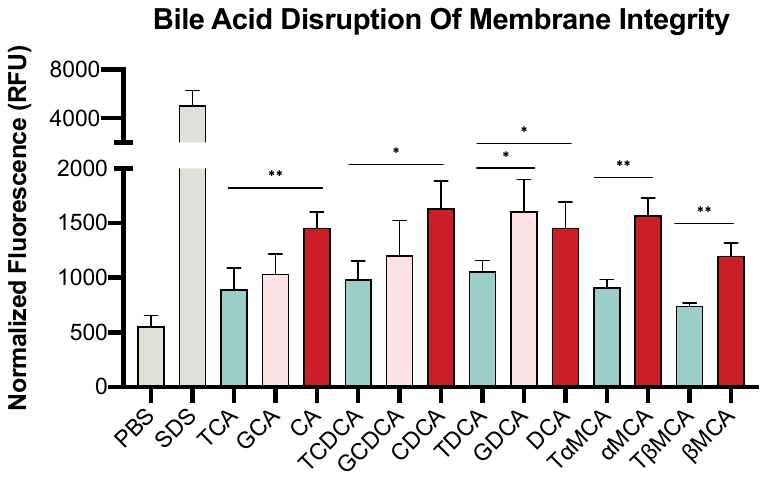

**Figure S5**

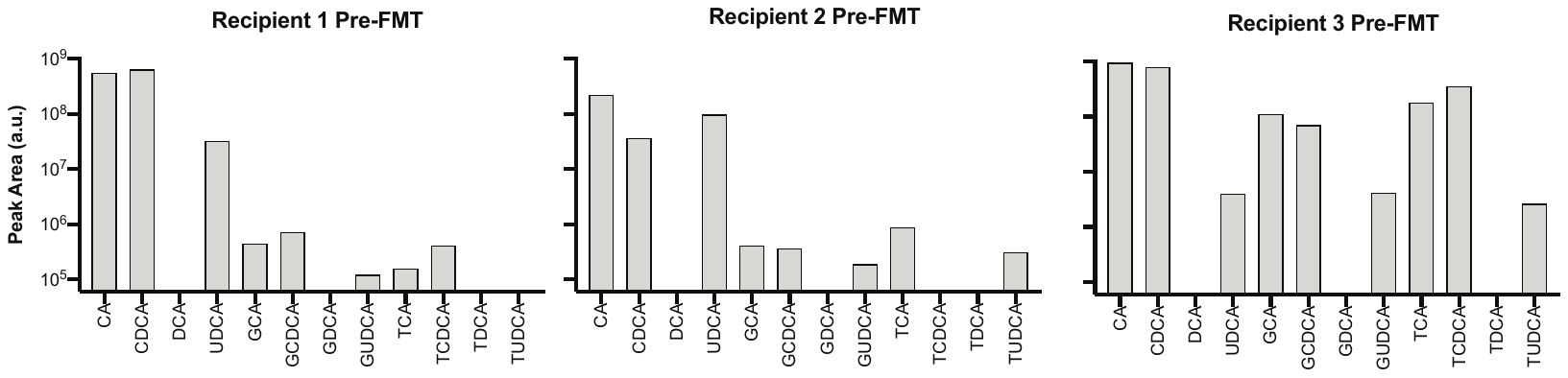

**Figure S6**

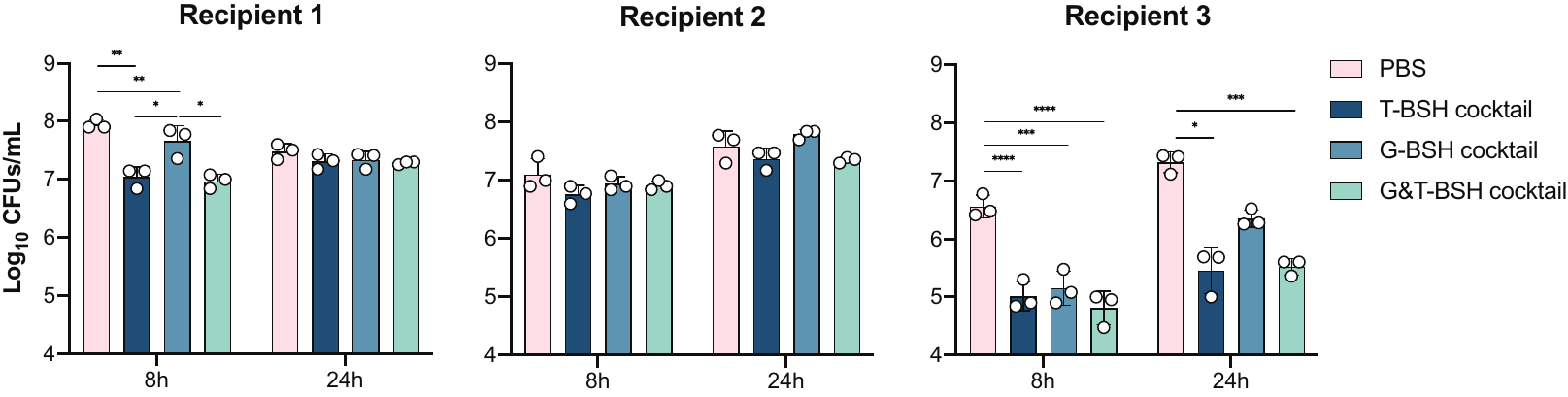

**Figure S7**

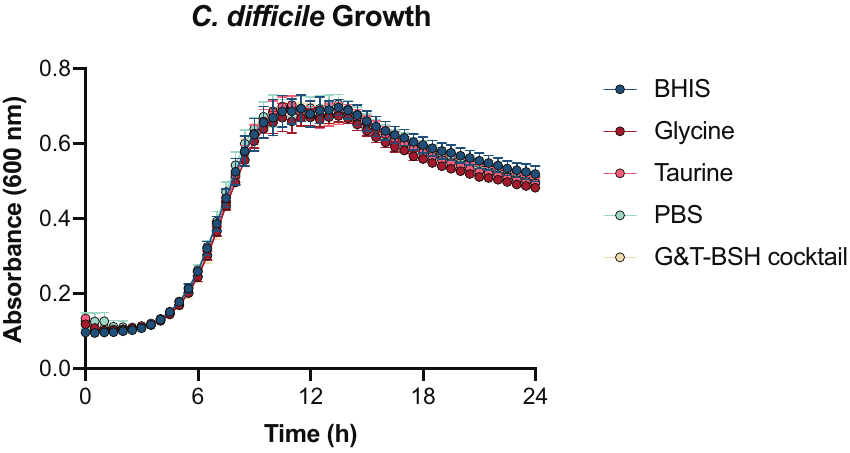

**Figure S8**

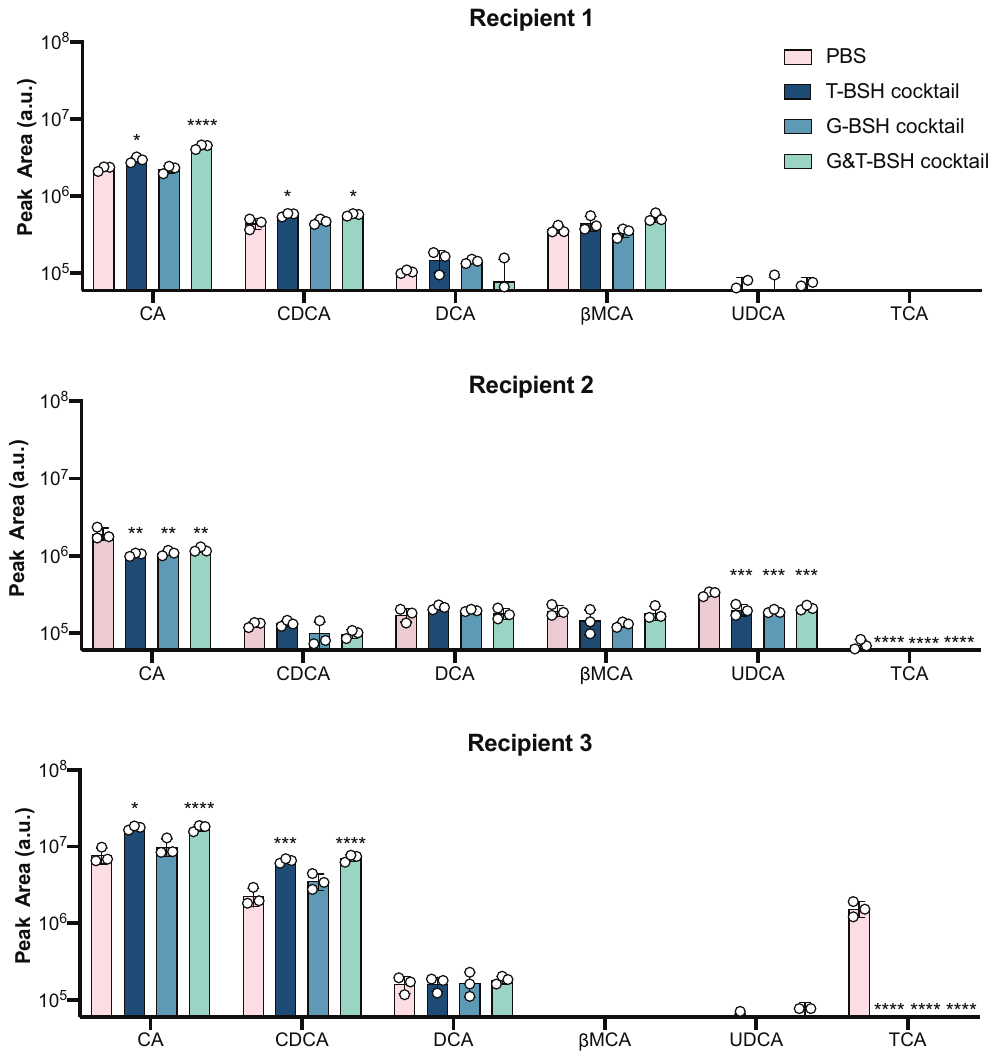

**Figure S9**

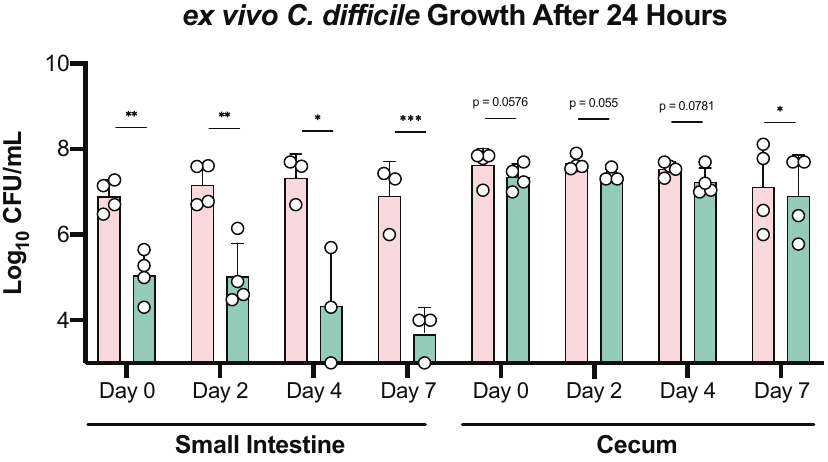

**Figure S10**

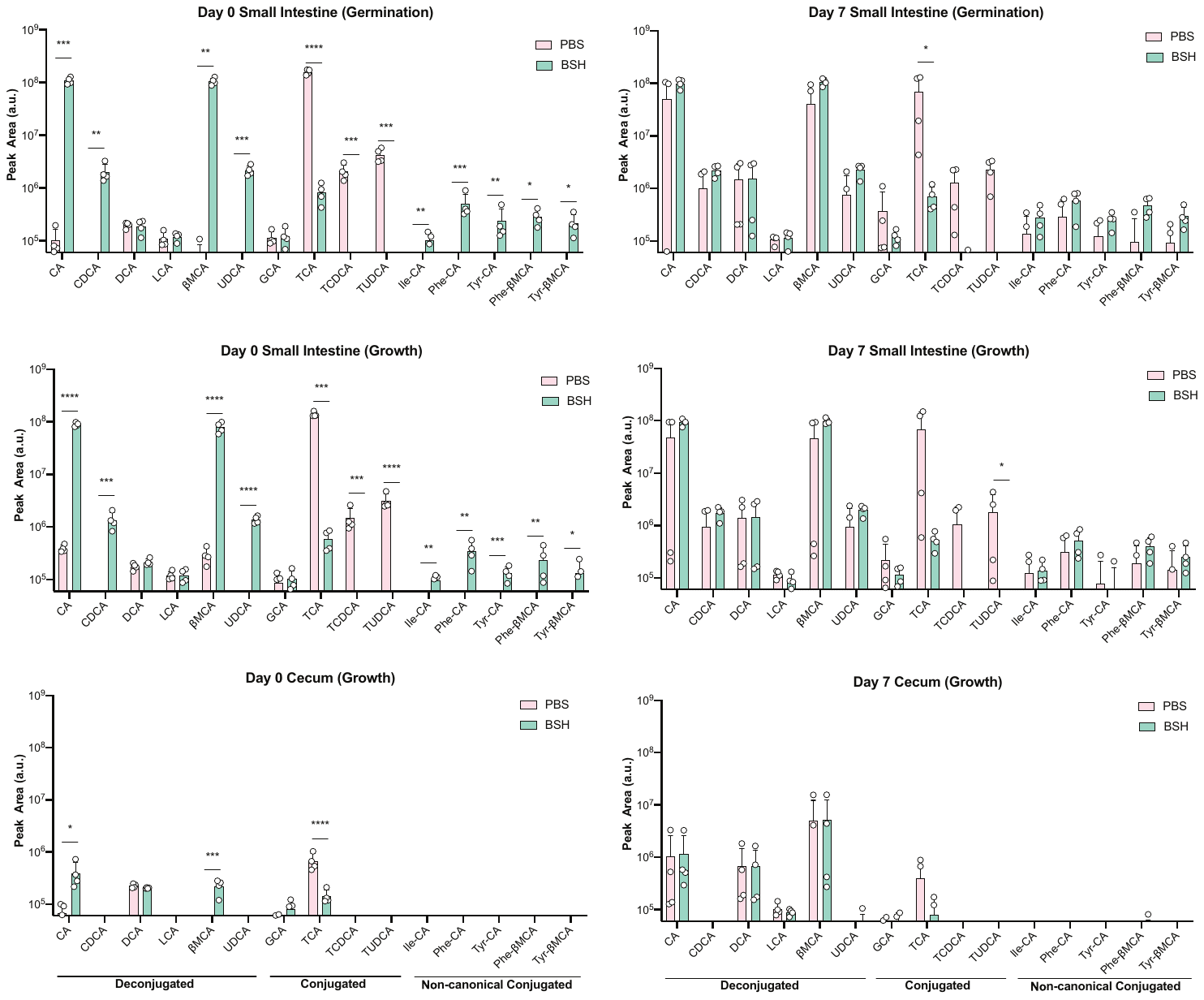

**Figure S11**

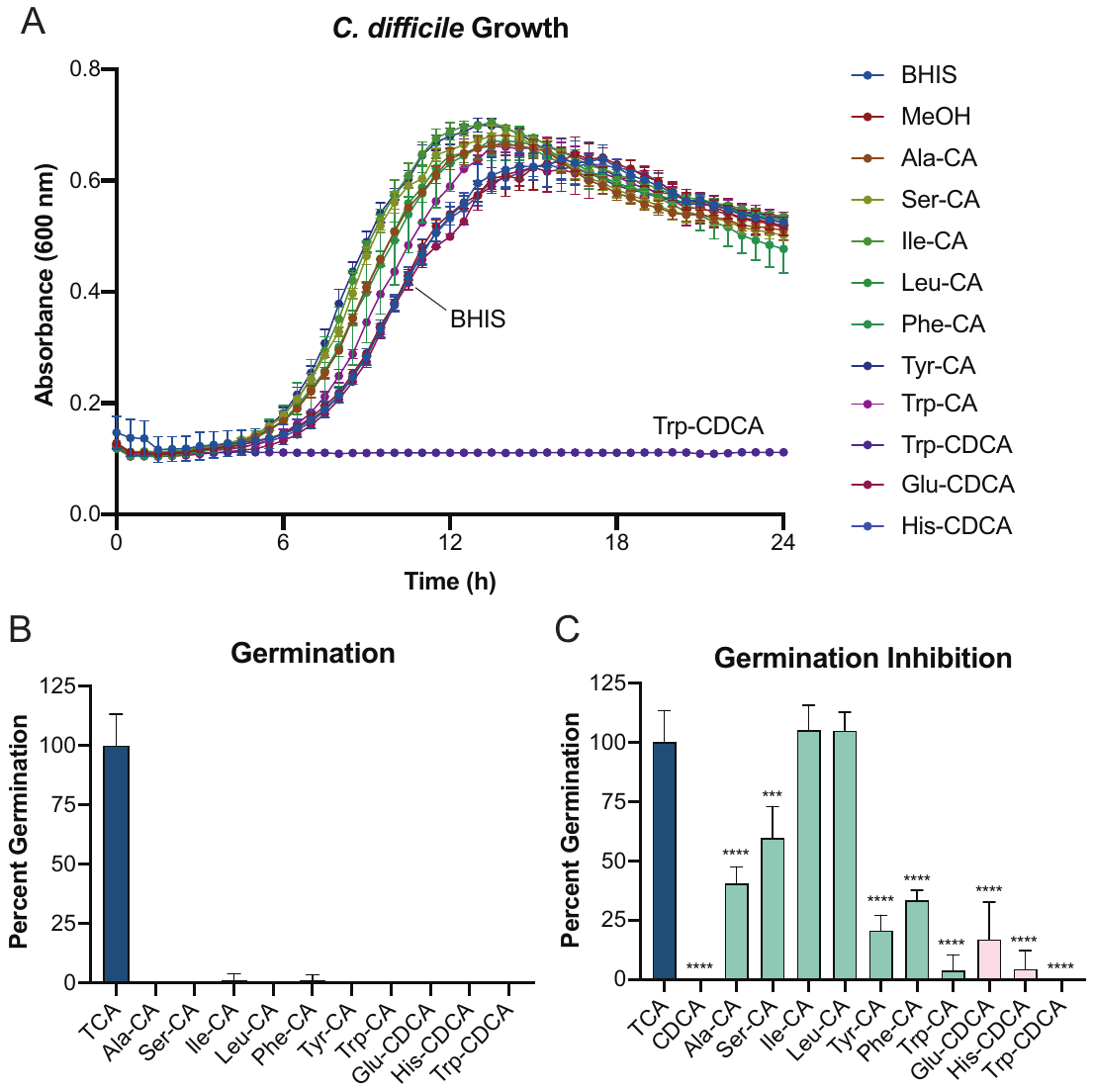

**Figure S12**

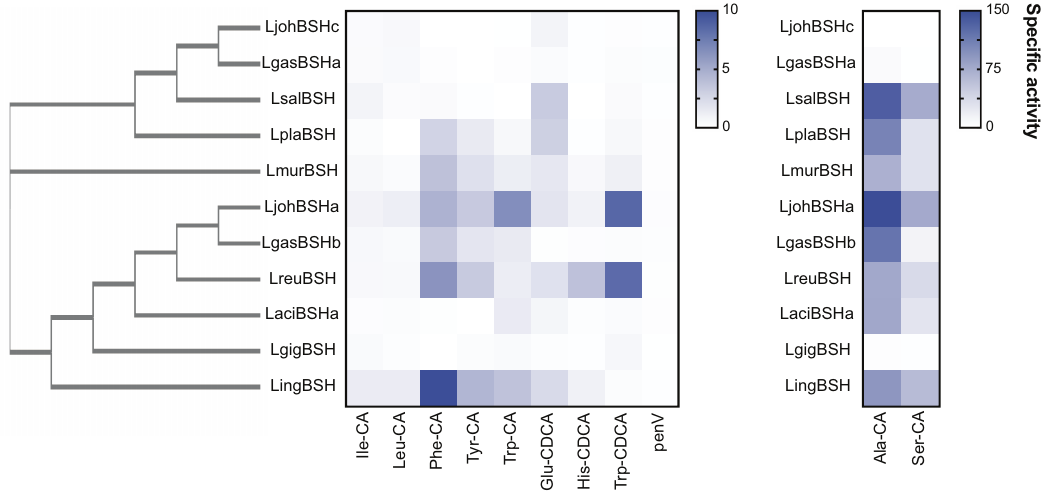

**Figure S13**

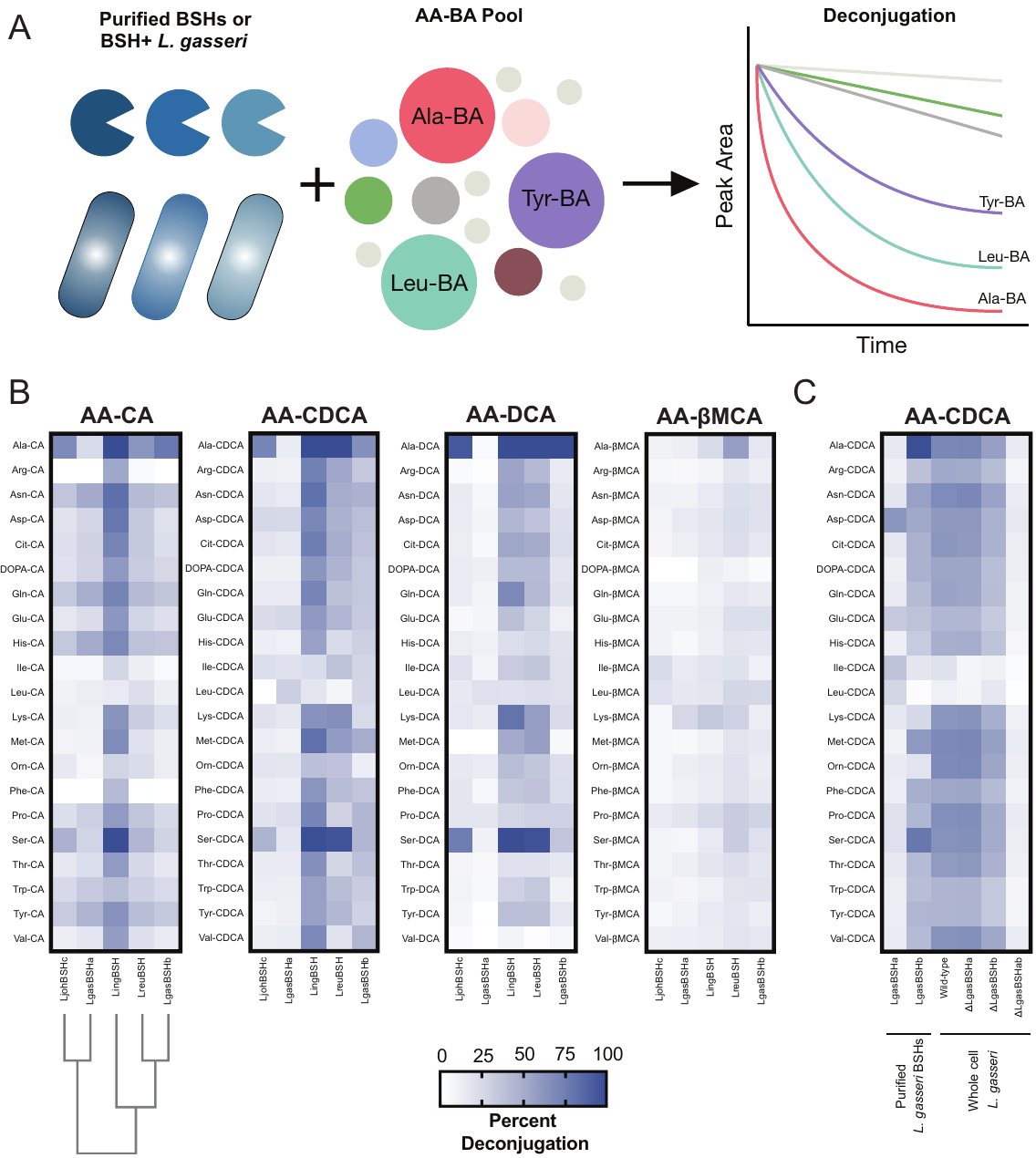

**Figure S14**

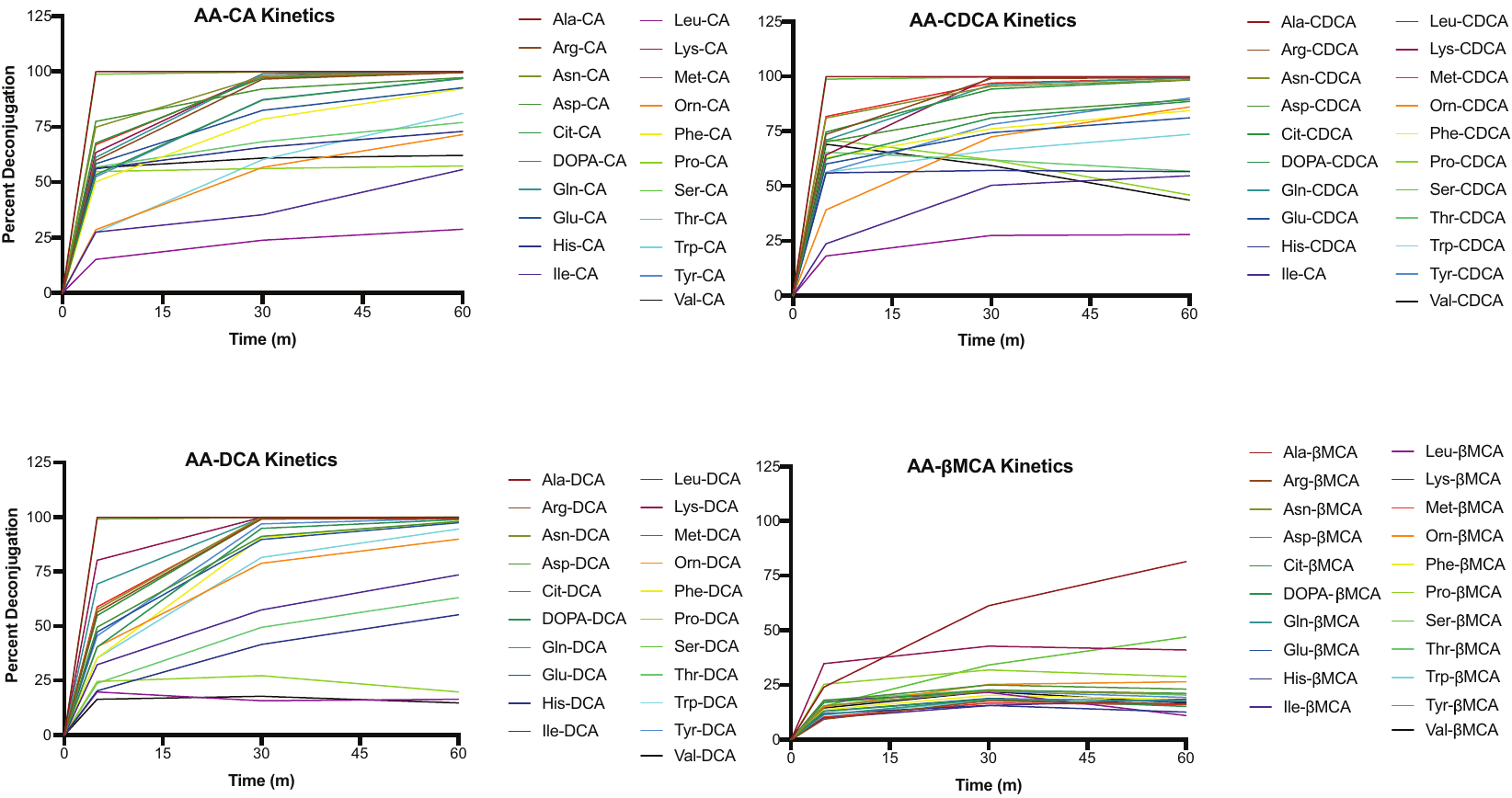

**Figure S15**

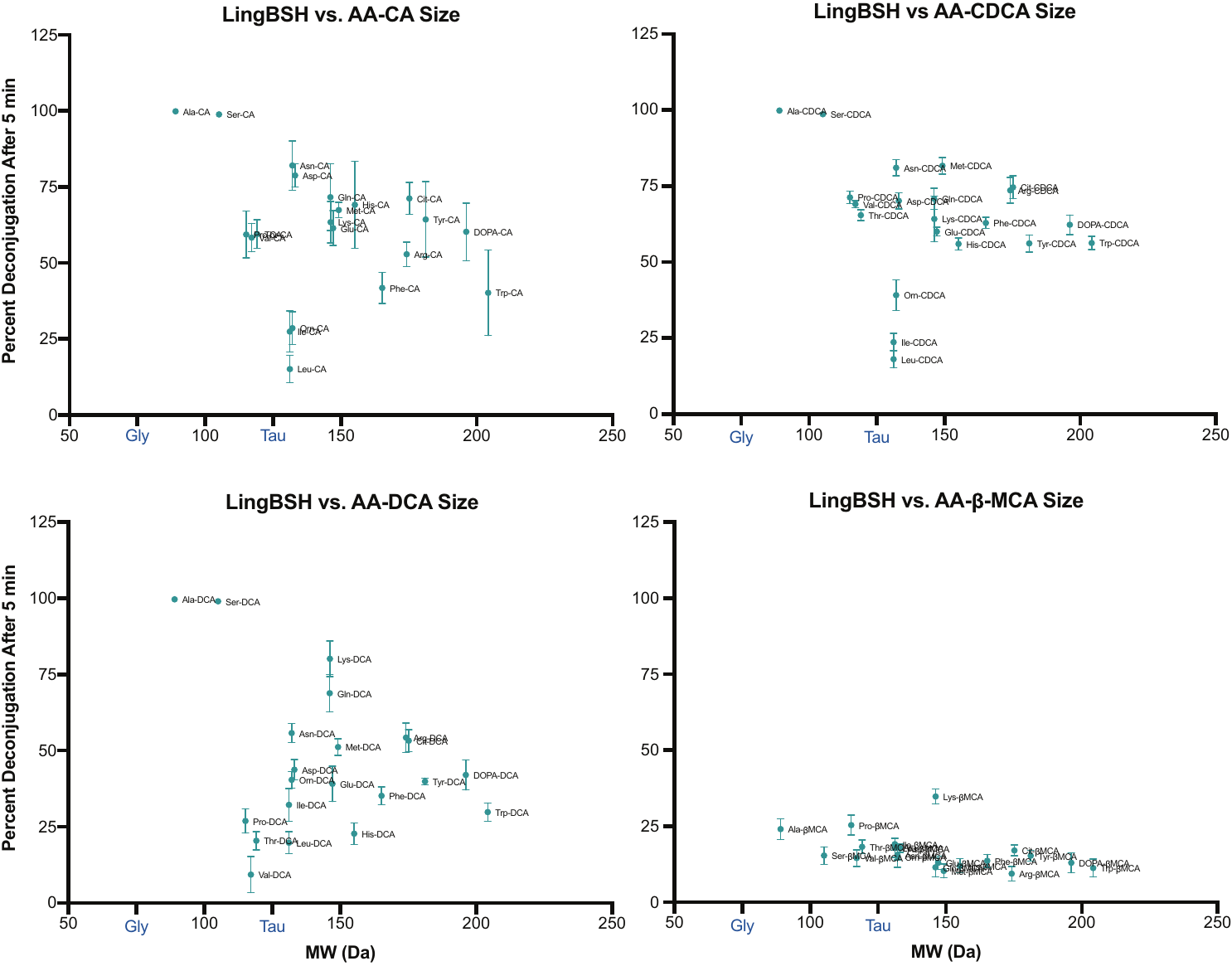

**Figure S16**

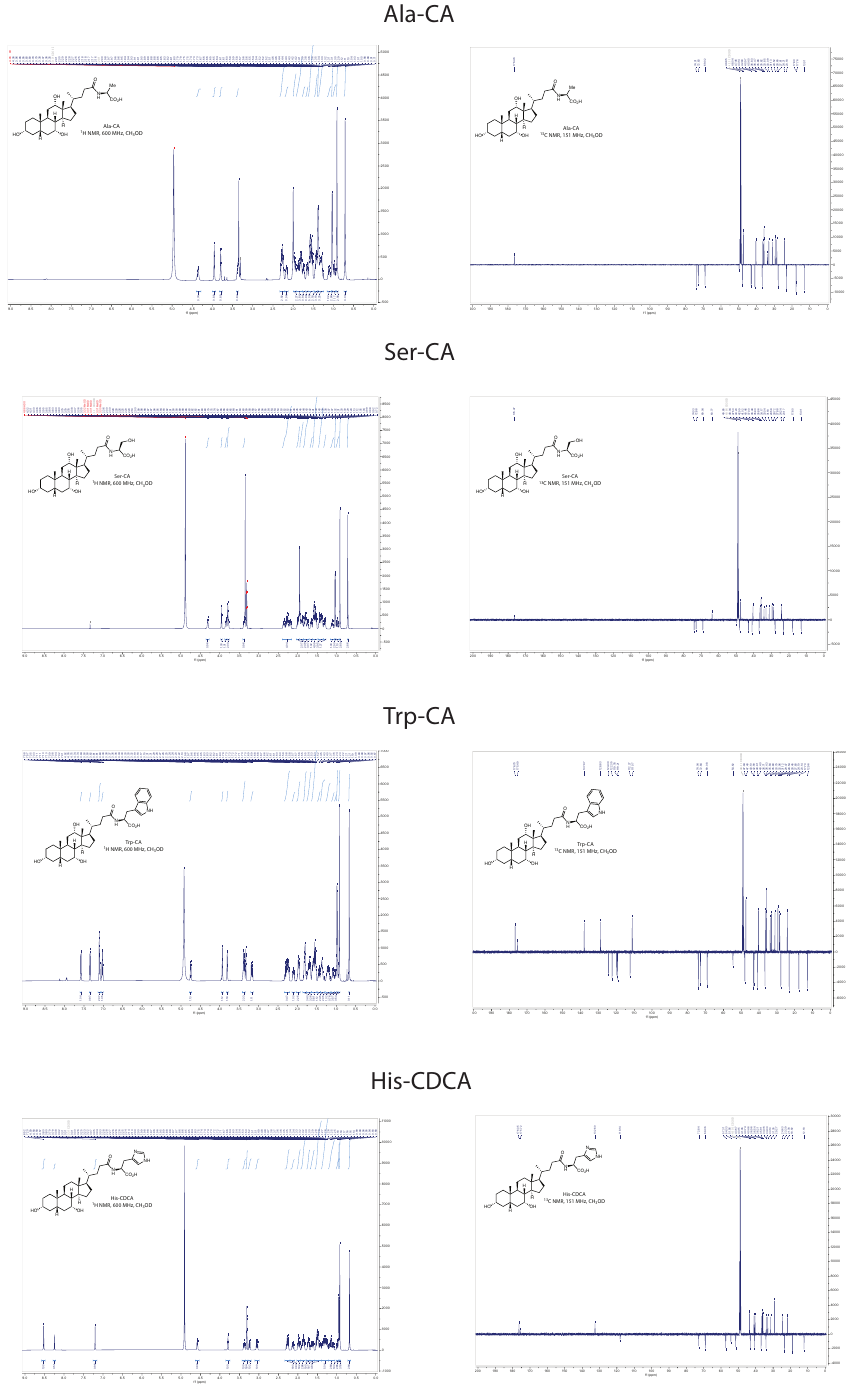

**Figure S17**

**Supplementary tables:**

**Table S1. *Lactobacillus* BSH cluster and phylogeny information**

**Table S2. Purified BSH sequences and information**

**Table S3. X-ray crystallography statistics**

**Table S1**

**Table S1 (continued)**

**Table S1 (continued)**

**Table S2**

**Table S3**
